# Highly variable molecular signatures of TDP-43 loss of function are associated with nuclear pore complex injury in a population study of sporadic ALS patient iPSNs

**DOI:** 10.1101/2023.12.12.571299

**Authors:** Jeffrey D. Rothstein, Caroline Warlick, Alyssa N. Coyne

## Abstract

The nuclear depletion and cytoplasmic aggregation of the RNA binding protein TDP-43 is widely considered a pathological hallmark of Amyotrophic Lateral Sclerosis (ALS) and related neurodegenerative diseases. Recent studies have artificially reduced TDP-43 in wildtype human neurons to replicate loss of function associated events. Although this prior work has defined a number of gene expression and mRNA splicing changes that occur in a TDP-43 dependent manner, it is unclear how these alterations relate to authentic ALS where TDP-43 is not depleted from the cell but miscompartmentalized to variable extents. Here, in this population study, we generate ∼30,000 qRT-PCR data points spanning 20 genes in induced pluripotent stem cell (iPSC) derived neurons (iPSNs) from >150 control, C9orf72 ALS/FTD, and sALS patients to examine molecular signatures of TDP-43 dysfunction. This data set defines a time dependent and variable profile of individual molecular hallmarks of TDP-43 loss of function within and amongst individual patient lines. Importantly, nearly identical changes are observed in postmortem CNS tissues obtained from a subset of patients whose iPSNs were examined. Notably, these studies provide evidence that induction of nuclear pore complex (NPC) injury via reduction of the transmembrane Nup POM121 in wildtype iPSNs is sufficient to phenocopy disease associated signatured of TDP-43 loss of function thereby directly linking NPC integrity to TDP-43 loss of function. Therapeutically, we demonstrate that the expression of *all* mRNA species associated with TDP-43 loss of function can be restored in sALS iPSNs via two independent methods to repair NPC injury. Collectively, this data 1) represents a substantial resource for the community to examine TDP-43 loss of function events in authentic sALS patient iPSNs, 2) demonstrates that patient derived iPSNs can accurately reflect actual TDP-43 associated alterations in patient brain, and 3) that targeting NPC injury events can be preclinically and reliably accomplished in an iPSN based platform of a sporadic disease.

## Introduction

Amyotrophic Lateral Sclerosis (ALS) is a fatal adult onset neurodegenerative disease that affects the motor circuitry (motor neurons, interneurons, glial cells) within the motor cortex and spinal cord of the CNS. About 10% of ALS cases are inherited and referred to as familial ALS (fALS). The remaining 90% of ALS cases are sporadic (sALS). Mutations in >20 genes with functions in multiple cellular pathways including proteostasis and RNA metabolism have been identified as causative of ALS. Although some of these genetic mutations, most notably the C9orf72 mutation, have been associated with both fALS and sALS, the vast majority of sALS cases occur with no currently known genetic cause ^1–6^.

ALS, like Frontotemporal Dementia (FTD) and Alzheimer’s Disease (AD), is considered a TDP-43 proteinopathy. At end-stage disease, the normally predominantly nuclear RNA binding protein TDP-43 is heterogeneously depleted from the nucleus of CNS cells and in a small subset of cells aggregates in the cytoplasm ^7–16^. Nuclear depletion, and in some cells complete nuclear clearance, is thought to precede cytoplasmic mislocalization ^17^ and has been demonstrated to lead to a loss of nuclear function ^18–27^. Although early studies evaluated alterations in gene expression and mRNA splicing that occur in mice following TDP-43 depletion ^24,26^, there is great dissimilarity between mouse and human TDP-43 targets due to a lack of conservation in the consensus sequence for TDP-43 binding in TDP-43 targets in mice ^19,23,28^. To more accurately evaluate the role of TDP-43 nuclear depletion in human disease, recent studies utilized induced pluripotent stem cell (iPSC) technologies to identify alterations in gene expression and mRNA splicing that occur upon artificial depletion of TDP-43 in human neurons ^19,20,25^. Critically, these studies have identified hundreds of human specific TDP-43 mRNA targets dysregulated at the level of gene expression or splicing. Although, some of these TDP-43 loss of function associated cryptic exon containing mRNAs have been detected in a small number of patient biofluids or postmortem tissues, variability in the magnitude of abundance was observed across individual patients ^23,25,27^. Further, the profile of dysregulation of multiple TDP-43 targets in authentic iPSN models and how this human *in vitro* model system reflects alterations in actual ALS patient CNS tissues remains unknown. The establishment of a preclinical model that faithfully reproduces TDP-43 loss of function in authentic sALS disease is essential for understanding the pathobiology of authentic sALS along with preclinical testing of new therapeutic strategies that may impact TDP-43 function.

In this resource, we utilize a qRT-PCR based panel comprised of 18 gene expression and splicing changes associated with TDP-43 loss of function as well as 2 negative controls to examine time dependent alterations in TDP-43 function in >150 different patient iPSC lines obtained from the Answer ALS program ^29^. We determined that the emergence of “molecular signatures of TDP-43 dysfunction” are time and differentiation protocol dependent in C9orf72 and sALS patient iPSNs. Importantly, there is extensive variability in the magnitude of individual gene expression and splicing changes both within and amongst individual patient iPSC lines. Critically, we compared iPSNs and postmortem tissues obtained from the same patient and find that our iPSN model system mimics the TDP-43 dysfunction signatures observed in postmortem CNS tissues. Thus, this highlights the utility of our *in vitro* model system of sALS disease for preclinical therapeutic testing. To this end, we present two strategies to alleviate nuclear pore complex injury that restore TDP-43 functionality in a population of sALS iPSNs. Collectively, this study represents a critical resource establishing authentic C9orf72 and sALS patient iPSNs as a preclinical model system for investigating mechanisms underlying TDP-43 dysfunction and examining therapeutic strategies for restoring TDP-43 function in ALS.

## Results

### Molecular hallmarks of TDP-43 loss of function are time dependent and variable within and amongst sALS patient iPSNs

Loss of nuclear TDP-43 function associated with histologic depletion of nuclear TDP-43 is considered an early pathophysiologic hallmark of ALS and related neurodegenerative diseases ^18–27^. In an effort to understand how loss of nuclear TDP-43 function contributes to disease pathogenesis, a number of recent studies have artificially knocked down TDP-43 in human neurons and in doing so identified hundreds of TDP-43 loss of function associated changes in gene expression and mRNA splicing ^19,20,25^. However, how these molecular changes in knockdown based systems relate to authentic ALS patient neurons remains unknown. To address this, we selected a targeted panel of 6 gene expression changes (*ELAVL3*, *PFKP*, *RCAN1*, *SELPLG*, *STMN2*, and *UNC13A*) and 12 splicing alterations associated with TDP-43 loss of function. We elected to focus on cryptic exon inclusion splicing events in *ACTL6B*, *ARHGAP32*, *CAMK2B*, *CDK7*, *DNM1*, *HDGFL2*, *MYO18A*, *NUP188*, *POLDIP3*, *SYT7*, *STMN2*, and *UNC13A*. We also included 2 negative control genes (*ACTIN* and *POM121*) not known to be regulated by TDP-43. In total, we examined these 20 mRNA species by qRT-PCR in 33 control, 12 C9orf72, and 111 sALS patient iPSC lines at 3 time points (day 32, 46, and 60) in our recently modified and optimized small molecule/growth factor mediated spinal neuron differentiation protocol ^30^. To account for variability occurring across different differentiations of the same line, each iPSC line was differentiated a total of 3 times. Overall, we observed a significant decrease in *ELAVL3*, *PFKP*, *RCAN1*, *SELPLG*, and *STMN2* mRNA expression (**Figures 1-2, Supplemental Figure 1, Supplemental File 1**) and a significant increase in cryptic exon containing *ACTL6B, ARHGAP32*, *CAMK2B*, *CDK7*, *DNM1*, *HDGFL2*, *MYO18A*, *SYT7*, and truncated *STMN2* mRNA expression at day 46 and 60, but not day 32 of differentiation (**Figures 3-4, Supplemental Figure 2, Supplemental File 1**). The expression of *ACTIN* and *POM121* remain unchanged. Interestingly, we were unable to detect changes in *UNC13A* (**Figures 1-2, Supplemental Figure 1, Supplemental File 1**) and cryptic exon containing *NUP188*, *POLDIP3*, and *UNC13A* (**Figures 3-4, Supplemental Figure 2, Supplemental File 1**) mRNA expression at the time points evaluated. Due to technical limitations, we were unable to examine time points beyond 60 days at this scale in our iPSN model. Longitudinal analysis suggests that within individual patient iPSC lines, the magnitude of TDP-43 loss of function associated alterations in gene expression and splicing are largely stable over time after emergence (**Figure 2**, **Figure 4, Supplemental Figure 1-2, Supplemental File 1**). However, for many patient lines, the magnitude of individual mRNA alterations increased over time and for a small subset some displayed a slight restoration back towards control levels (**Figure 2**, **Figure 4, Supplemental Figure 1-2, Supplemental File 1**). Perhaps most strikingly, we observed extensive heterogeneity in the magnitude of individual mRNA dysregulation events amongst individual patients (**Figures 1-4, Supplemental Figures 1-2, Supplemental File 1**). Moreover, within individual patients there was a widespread discordance in the magnitude of individual mRNA dysregulation events (**Figures 1-4, Supplemental Figures 1-2, Supplemental File 1**). For example, *STMN2*, originally identified as one of the most altered mRNA targets following artificial TDP-43 knockdown in SH-SY5Y cells and iPSNs ^19,23^, was variably altered in this large population of patient iPSNs, with a substantial number of patient showing only modest changes in *STMN2* expression and splicint, with other TDP-43 loss of function associated mRNA changes in some cases more dramatically altered (e.g. *ELAVL3*, *MYO18A* CE) (**Figures 1-4, Supplemental Figures 1-2, Supplemental File 1**). When taken as an aggregate “TDP-43 dysfunction score”, the variability of overall magnitude of TDP-43 associated loss of function amongst individual patient iPSNs remained (**Supplemental Figure 3a**). However, for a subset of patient lines where TDP-43 localization was monitored by immunostaining and confocal imaging, there was a correlation between nuclear/cytoplasmic distribution of TDP-43 and TDP-43 dysfunction such that a decrease in the nuclear/cytoplasmic ratio of TDP-43 immunoreactivity correlated with increased overall dysfunction over time (**Supplemental Figure 3b-d**). We have previously demonstrated that decreased nuclear/cytoplasmic TDP-43 ratios occur in the absence of TDP-43 aggregation in our iPSN model ^30,31^. Eliminating control iPSNs from this analysis lessened the overall correlation between TDP-43 localization and dysfunction, albeit a subtly significant correlation remained at day 60 of differentiation (**Supplementary Figure 3e-g**). Molecular hallmarks of TDP-43 dysfunction were not detected in mutant SOD1 patient iPSNs at day 32, 46 (**Supplemental File 1**) or 60 (**Supplemental Figure 4, Supplemental File 1**) of differentiation or in postmortem patient mutant SOD1 CNS tissues (**Supplemental Figure 5**) consistent with the lack of TDP-43 pathology in SOD1 ALS ^32,33^. On the other hand, a small decrease in *STMN2* mRNA and small increase in truncated *STMN2* mRNA were observed at day 60 (**Supplemental Figure 6, Supplemental File 1**), but not day 32 or 46 (**Supplemental File 1**) of differentiation in mutant TDP-43 patient iPSNs. No other TDP-43 loss of function associated gene expression or mRNA splicing alterations were observed in mutant TDP-43 patient iPSNs at the time points evaluated (**Supplemental Figure 6, Supplemental File 1**). Collectively, our data indicates that the emergence of molecular signatures of TDP-43 dysfunction are time dependent and heterogenous within and amongst authentic ALS patient iPSNs, with the most prominent changes seen in the large population of sALS lines.

**Figure 1:**
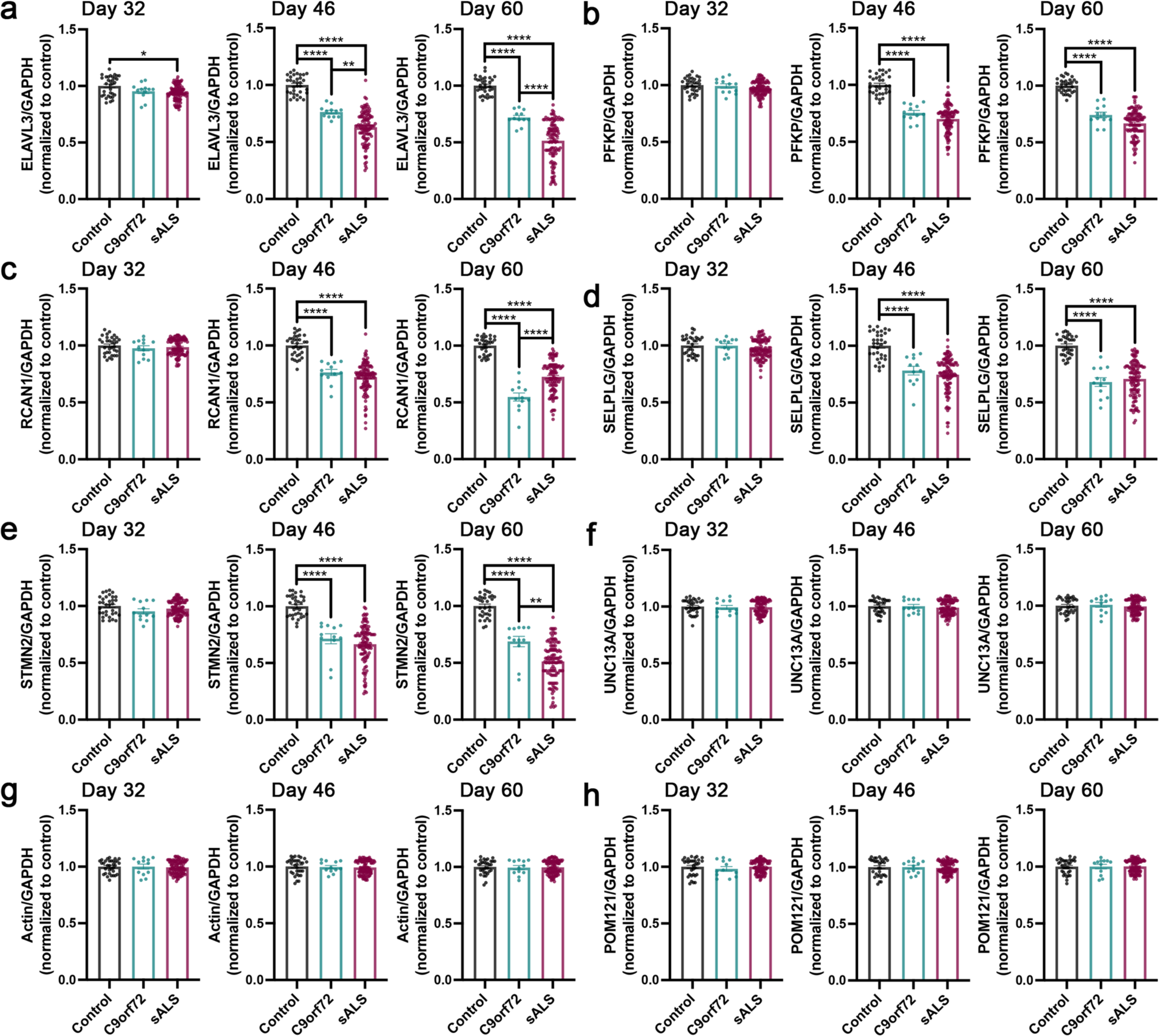
TDP-43 loss of function associated changes in target gene expression are time dependent in C9orf72 ALS/FTD and sALS iPSNs. **(a-h)** qRT-PCR for *ELAVL3* **(a)**, *PFKP* **(b)**, *RCAN1* **(c)**, and *SELPLG* **(d)**, *STMN2* **(e)**, *UNC13A* **(f)**, *ACTIN* **(g)**, and *POM121* **(h)** mRNA in control, C9orf72, and sALS iPSNs at day 32, 46, and 60 of differentiation. GAPDH was used as the reference gene for normalization. *ACTIN* and *POM121* were used as negative control mRNAs not known to be regulated by TDP-43. n = 33 control, 12 C9orf72, and 111 sALS iPSC lines. Data points for each line represent the average value across 3 independent differentiations. One-way ANOVA with Tukey’s multiple comparison test was used to calculate statistical significance. * p < 0.05, ** p < 0.01, **** p < 0.0001.

**Figure 2:**
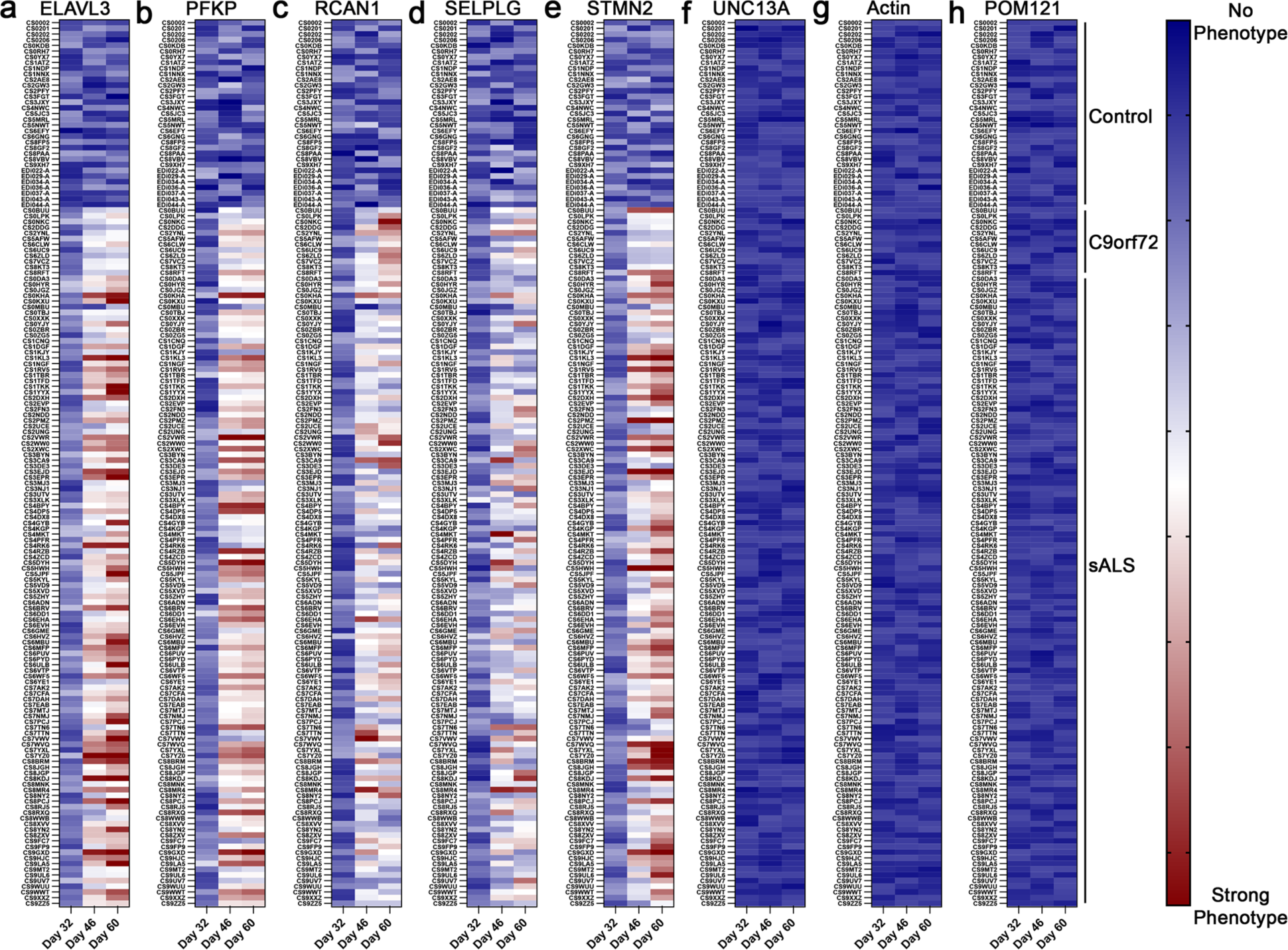
Time dependent TDP-43 loss of function associated changes in target gene expression are variable within and amongst individual C9orf72 ALS/FTD and sALS patient iPSNs. (**a-h**) Heat maps from qRT-PCR for *ELAVL3* (**a**), *PFKP* (**b**), *RCAN1* (**c**), and *SELPLG* (**d**), *STMN2* (**e**), *UNC13A* (**f**), *ACTIN* (**g**), and *POM121* (**h**) mRNA in control, C9orf72, and sALS iPSNs at day 32, 46, and 60 of differentiation. *ACTIN* and *POM121* represent negative control mRNAs not known to be regulated by TDP-43. n = 33 control, 12 C9orf72, and 111 sALS iPSC lines. For each line and time point, boxes represent the average value across 3 independent differentiations.

**Figure 3:**
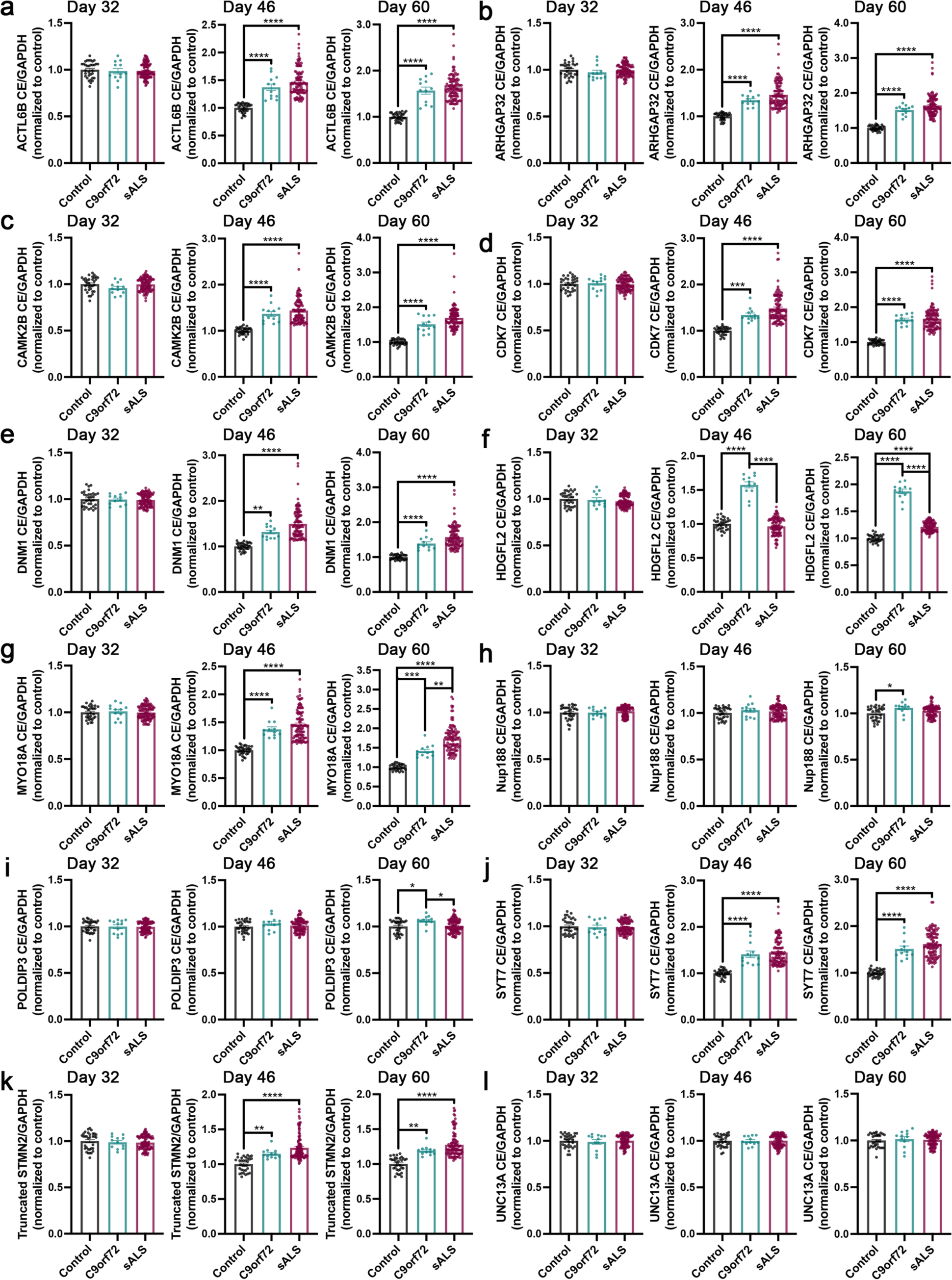
TDP-43 loss of function associated alterations in target mRNA splicing are time dependent in C9orf72 ALS/FTD and sALS iPSNs. (**a-l**) qRT-PCR for *ACTL6B* cryptic exon containing (**a**), *ARHGAP32* cryptic exon containing (**b**), *CAMK2B* cryptic exon containing (**c**), *CDK7* cryptic exon containing (**d**), *DNM1* cryptic exon containing (**e**), *HDGFL2* cryptic exon containing (**f**), *MYO18A* cryptic exon containing (**g**), *NUP188* cryptic exon containing (**h**), *POLDIP3* cryptic exon containing (**i**), *SYT7* cryptic exon containing (**j**), truncated *STMN2* (**k**), and *UNC13A* cryptic exon containing (**l**) mRNA in control, C9orf72, and sALS iPSNs at day 32, 46, and 60 of differentiation. GAPDH was used as the reference gene for normalization. n = 33 control, 12 C9orf72, and 111 sALS iPSC lines. Data points for each line represent the average value across 3 independent differentiations. One-way ANOVA with Tukey’s multiple comparison test was used to calculate statistical significance. * p < 0.05, ** p < 0.01, *** p < 0.001, **** p < 0.0001.

**Figure 4:**
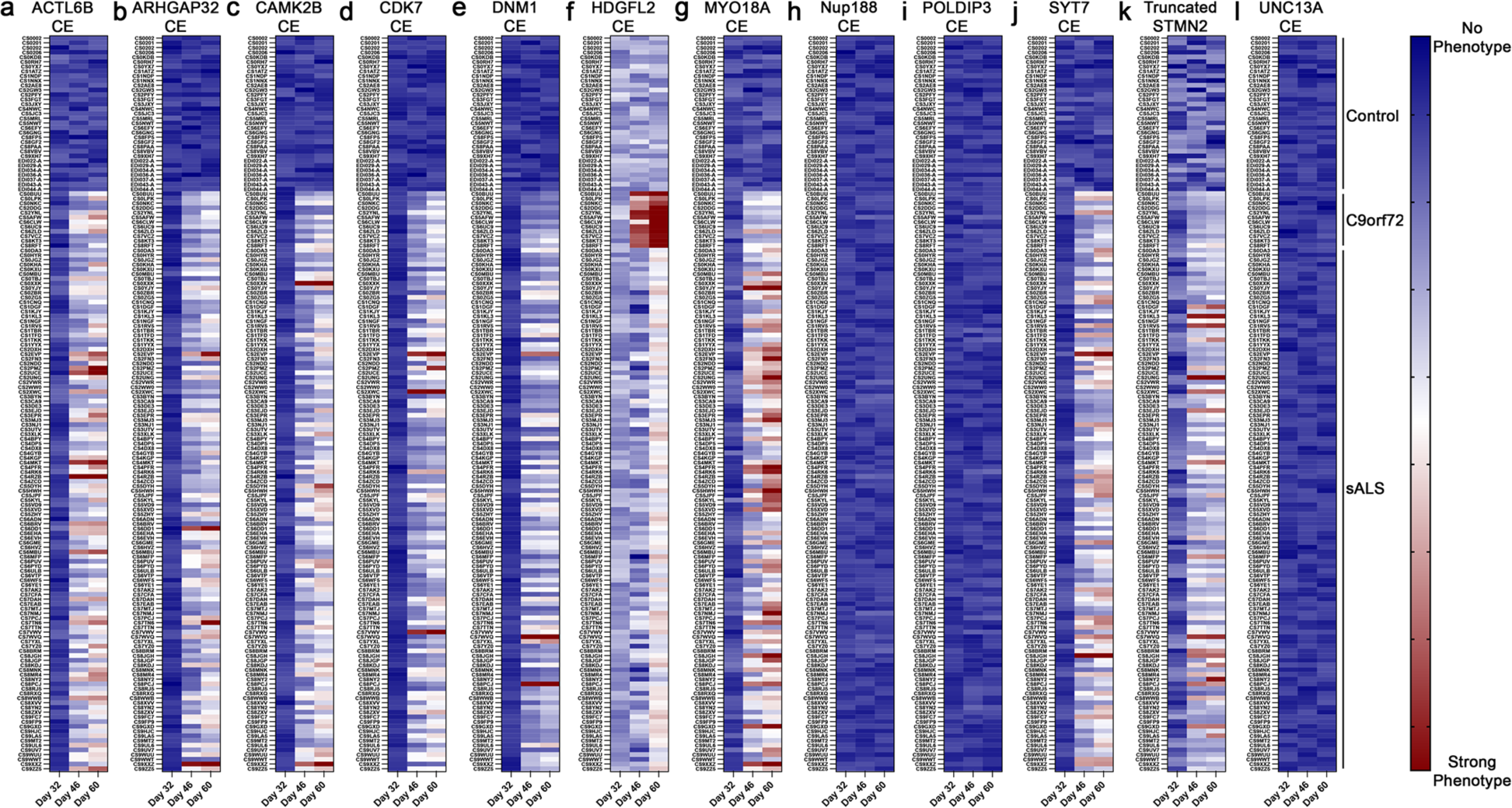
Time dependent TDP-43 loss of function associated alterations in target mRNA splicing are variable within and amongst individual C9orf72 ALS/FTD and sALS patient iPSNs. (**a-l**) Heat maps from qRT-PCR for *ACTL6B* cryptic exon containing (**a**), *ARHGAP32* cryptic exon containing (**b**), *CAMK2B* cryptic exon containing (**c**), *CDK7* cryptic exon containing (**d**), *DNM1* cryptic exon containing (**e**), *HDGFL2* cryptic exon containing (**f**), *MYO18A* cryptic exon containing (**g**), *NUP188* cryptic exon containing (**h**), *POLDIP3* cryptic exon containing (**i**), *SYT7* cryptic exon containing (**j**), truncated *STMN2* (**k**), and *UNC13A* cryptic exon containing (**l**) mRNA in control, C9orf72, and sALS iPSNs at day 32, 46, and 60 of differentiation. n = 33 control, 12 C9orf72, and 111 sALS iPSC lines. For each line and time point, boxes represent the average value across 3 independent differentiations.

### Detection of molecular hallmarks of TDP-43 loss of function is dependent upon iPSN differentiation protocol

Due to the ease of use and generation of “pure” populations of a single cell type, a number of groups now employ inducible iPSC differentiation protocols. These protocols generate homogenous populations of “cortical-like” or lower motor neurons based on the doxycycline inducible expression of NGN2 or NGN2/ISL1/LHX3 transcription factors respectively ^34,35^ and resulting cells are commonly referred to as “iNeurons”. Our recent study suggests that the magnitude of individual gene expression and mRNA splicing alterations following artificial TDP-43 knockdown in our mixed spinal neuron cultures is distinct from those observed following TDP-43 knockdown in SH-SY5Y cells or homogenous “cortical-like” iPSC derived neuronal cultures generated via induction of transcription factor expression differentiation protocols ^19,20,23,25,30^. As a result, we wanted to compare our results from our authentic patient iPSNs generated with our spinal neuron differentiation protocol ^30^ to other commonly used iPSN differentiation methods. Additionally, given that ∼50% of ALS patients exhibit a mild cognitive decline, largely in executive functioning, and 15% are diagnosed with FTD predominantly prior to the onset of ALS symptoms ^5,36,37^, we also wanted to ask whether molecular signatures of TDP-43 dysfunction could also be observed in other neuronal populations, namely cortical neurons. Therefore, we selected a subset of 3 control and 6 sALS patient iPSC lines and differentiated them using 4 differentiation protocols. To examine variability arising from technical differentiation replicates, iPSC lines were differentiated with each protocol a total of 3 times. Compared to controls (horizontal dashed line represents normalized control values for each differentiation protocol), ALS iPSNs generated using small molecule/growth factor mediated cortical ^38^ and spinal ^30^ neuron differentiation protocols that reproduce human neurodevelopmental processes displayed a similar magnitude of TDP-43 dysfunction across all mRNA species evaluated at day 120 and day 60 of differentiation respectively (**Figure 5**). In contrast, we were unable to observe molecular hallmarks of TDP-43 dysfunction in 30 day old “cortical-like” and lower motor neurons (**Figure 5**) generated via inducible transcription factor mediated differentiation protocols ^34,35^ using piggybac technology ^39–42^. Notably, in our hands, these neurons (including those generated from control iPSC lines) visibly displayed signs of cellular stress/death such as neurite fragmentation reproducibly by day 35 in culture. Subsequently, all neurons reproducibly died by day 40 in culture, with some iPSC lines succumbing to death even at earlier time points. Taken together, these data support a time dependent emergence of detectable TDP-43 dysfunction in authentic patient iPSNs and additionally suggest that culture complexity may play a role in the development of TDP-43 loss of function in ALS iPSNs.

**Figure 5:**
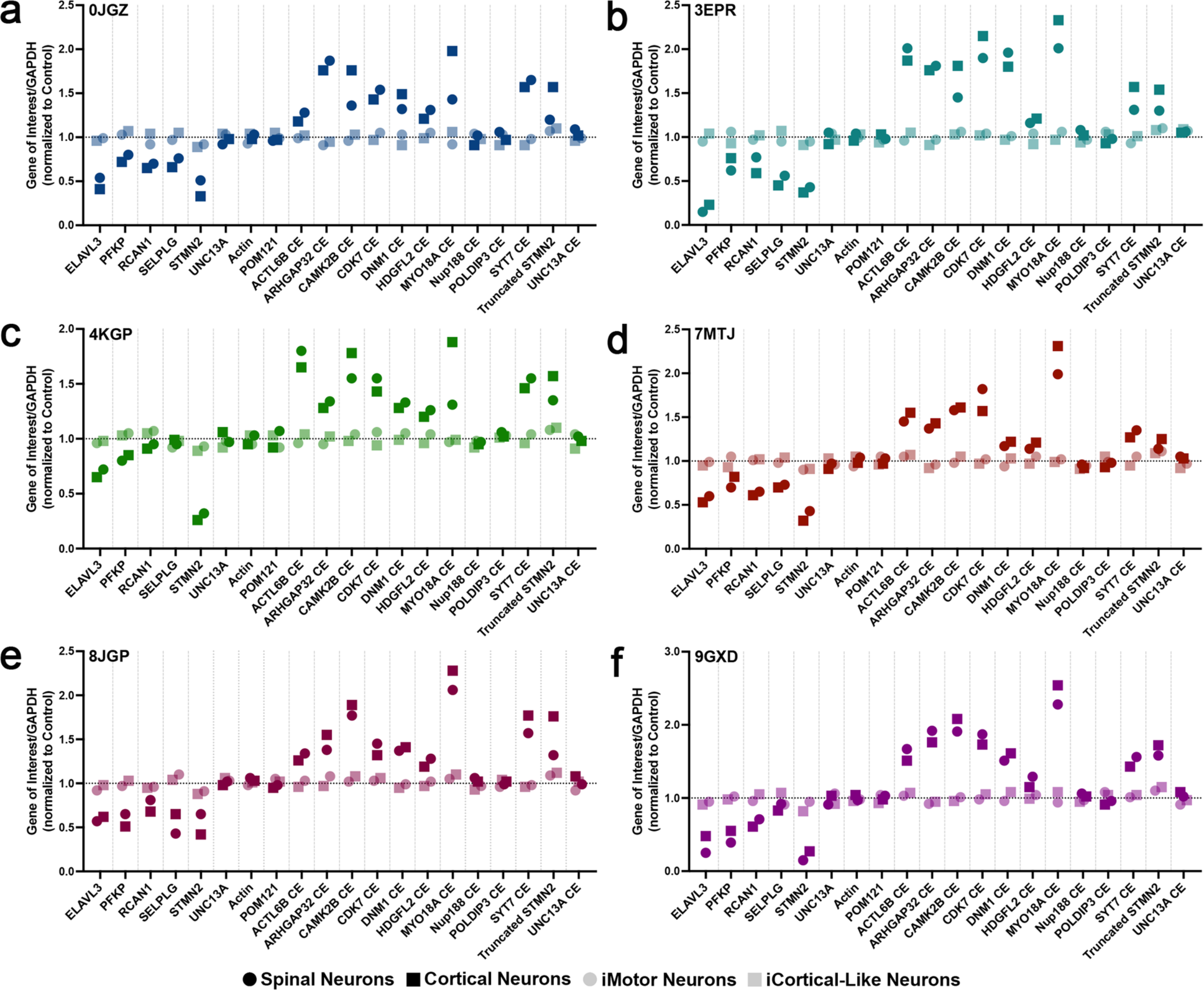
Molecular signatures of TDP-43 dysfunction are not observed in iPSNs generated with inducible transcription factor based differentiation protocols. (**a-f**) qRT-PCR for TDP-43 loss of function associated changes in gene expression and mRNA splicing in iPSC lines differentiated with 4 commonly utilized iPSN differentiation protocols. GAPDH was used as the reference gene for normalization. *ACTIN* and *POM121* were used as negative control mRNAs not known to be regulated by TDP-43. Deidentified iPSC line code as indicated in upper left. Symbols indicate differentiation protocol as detailed in legend. Each data point represents the average value across 3 independent differentiations per iPSC line and protocol. Normalized control values represented by horizontal dashed line. 6 age, sex, and passage matched control iPSC lines were used for iPSN analyses.

### Molecular hallmarks of TDP-43 loss of function in patient iPSNs mimic postmortem patient pathology

A critical component in determining the utility of preclinical models of disease is an evaluation of how closely they recapitulate real human disease pathophysiology. We have previously demonstrated that our iPSN model accurately reproduces alterations in the neuronal nuclear pore complex (NPC) that are commonly detected in actual sALS and C9orf72 CNS tissues^31,43^. As a result, we asked whether the iPSC derived spinal neuron model of ALS sufficiently recapitulated TDP-43 loss of function events detected in real human CNS. To this end, we obtained available postmortem human CNS tissues from 4 patients whose iPSNs we had previously examined (**Figures 1-4, Supplemental Figures 1-3, Supplemental File 1**) for TDP-43 loss of function associated mRNA misprocessing events. Employing a qRT-PCR based panel of gene expression and mRNA splicing alterations associated with TDP-43 dysfunction, we observed a strikingly similar magnitude of dysregulation of individual mRNA processing events in day 60 iPSC derived spinal neurons compared to cervical spinal cord and multiple motor cortex samples from the same patients (**Figure 6a-d**). We did not detect TDP-43 loss of function associated mRNA misprocessing events in the occipital cortex (**Figure 6a-d**) which is pathologically and clinically unaffected in ALS. In fact, with the exception of truncated *STMN2* and cryptic exon containing *UNC13A*, cryptic exon containing mRNAs were not detected in control (average value across 4 control cases represented by horizontal dashed line) or ALS postmortem occipital cortex (**Figure 6a-d**). Together, these data demonstrate that ALS iPSC derived spinal neuron models, such as that utilized in this study, accurately and faithfully reproduce molecular signatures of TDP-43 dysfunction detected in the same patient’s autopsied motor cortex and spinal cord tissues.

**Figure 6:**
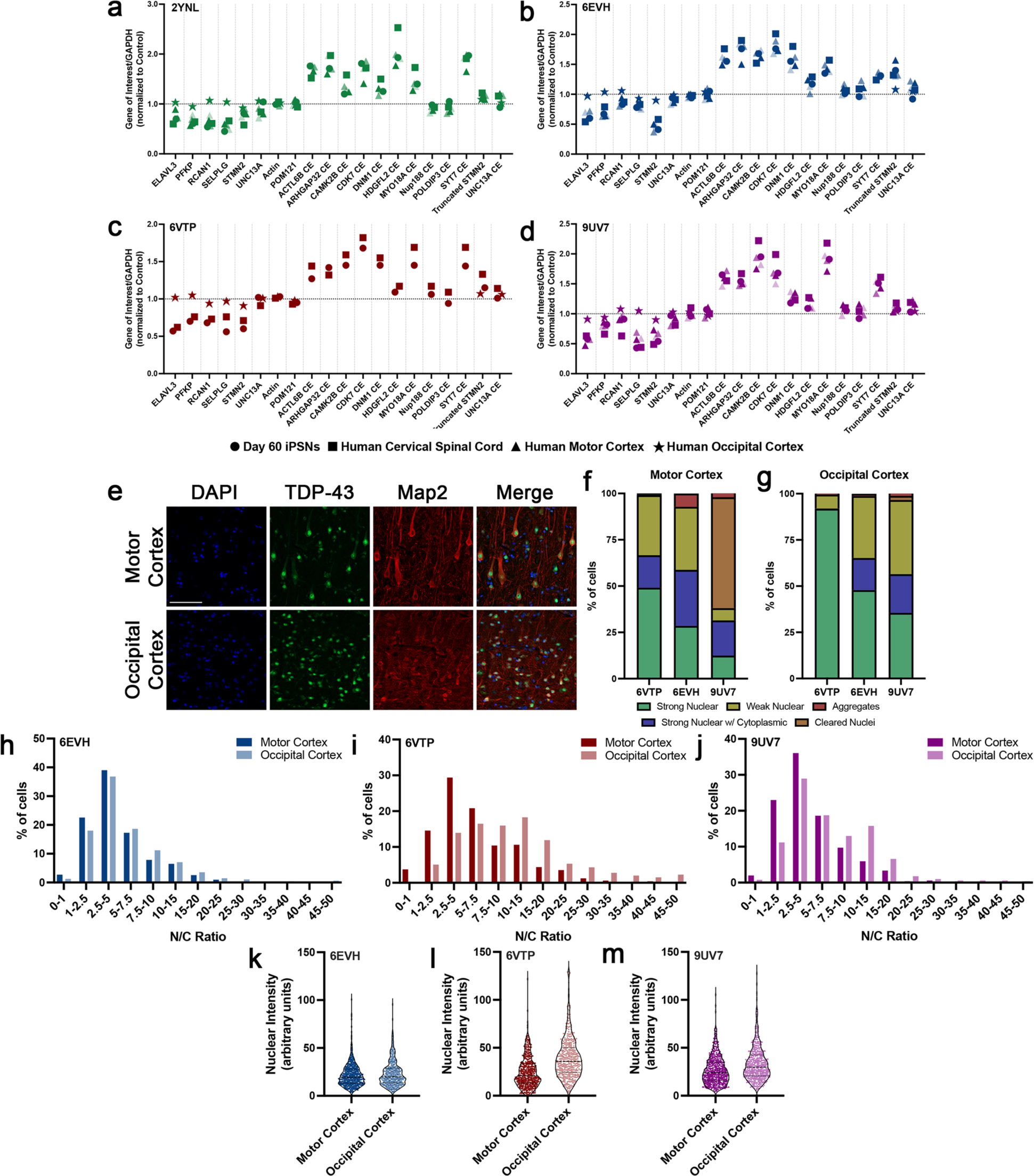
Molecular signatures of TDP-43 dysfunction in mixed spinal iPSN cultures are reminiscent of those observed in end-stage disease tissue and are independent of robust TDP-43 aggregation. (**a-d**) qRT-PCR for TDP-43 loss of function associated changes in gene expression and mRNA splicing in day 60 iPSNs compared to postmortem tissues from the same patients. GAPDH was used as the reference gene for normalization. *ACTIN* and *POM121* were used as negative control mRNAs not known to be regulated by TDP-43. Deidentified patient code as indicated in upper left. Symbols represent sample type as detailed in legend. iPSN data points represent the average value from 3 independent differentiations. Normalized control values represented by horizontal dashed line. 4 non-neurological controls were used for postmortem human tissue analyses. 4 age, sex, and passage matched control iPSC lines were used for iPSN analyses. (**e**) Immunostaining and apotome based imaging of TDP-43 in postmortem human tissues. Antibody for immunostaining as indicated on top, brain region as indicated on left. Scale bar = 50 μm. (**f-g**) Quantification of percentage of cells displaying various TDP-43 subcellular distribution patterns in postmortem motor (**f**) and occipital (**g**) cortex. n = 3 sALS patients, 1509 cells (6EVH motor cortex), 1427 cells (6EVH occipital cortex), 1012 cells (6VTP motor cortex), 1141 cells (6VTP occipital cortex), 1133 cells (9UV7 motor cortex), 1775 cells (9UV7 occipital cortex). (**h-j**) Distribution of TDP-43 nuclear/cytoplasmic ratios in postmortem sALS motor and occipital cortex neurons. Deidentified sALS patient code as indicated in upper left. n = 510 neurons (6EVH motor cortex), 484 neurons (6EVH occipital cortex), 480 neurons (6VTP motor cortex), 394 neurons (6VTP occipital cortex), 505 neurons (9UV7 motor cortex), 501 neurons (9UV7 occipital cortex). (**k-m**) TDP-43 nuclear intensity in postmortem sALS motor and occipital cortex neurons. Deidentified sALS patient code as indicated in upper left.

Next, similar to our iPSN studies (**Supplemental Figure 3**), we wanted to examine how TDP-43 dysfunction may relate to histologic TDP-43 pathology in end-stage CNS autopsy tissues. Therefore, we performed immunostaining and apotome based imaging for TDP-43 in postmortem motor and occipital cortex tissues (**Figure 6e**) from 3 of the 4 cases examined for TDP-43 loss of function events. Consistent with prior reports that TDP-43 aggregates are rare in ALS patient autopsy tissue ^14,15^, we found that only a small percentage of cells in the motor cortex contained cytoplasmic TDP-43 aggregates (**Figure 6f**). However, a number of cells within both the motor and occipital cortex displayed varying degrees of diffuse nuclear and cytoplasmic immunoreactivity (**Figure 6f-g**) albeit with an overall higher percentage of cells with robust nuclear TDP-43 immunoreactivity detected in the occipital cortex (**Figure 6g**). Notably, we only observed nuclei completely devoid of TDP-43 immunoreactivity in only one case (**Figure 6f-g**) with a higher percentage of neurons with nuclear TDP-43 clearance in motor (**Figure 6f**) compared to occipital (**Figure 6g**) cortex. Moreover, when we examined the distribution of nuclear/cytoplasmic ratios of TDP-43, we found an increased percentage of neurons with lower nuclear/cytoplasmic TDP-43 ratios in motor compared to occipital cortex in all 3 cases evaluated (**Figure 6h-j**) consistent with increased cytoplasmic immunoreactivity and/or decreased nuclear immunoreactivity. Moreover, nuclear intensity of TDP-43 immunostaining appeared to be slightly increased in neurons within the occipital cortex compared to motor cortex in 2 out of 3 cases examined (**Figure 6k-m**). Nuclear intensity moderately correlated with nuclear/cytoplasmic distribution of TDP-43 in postmortem motor and occipital cortex (**Supplemental Figure 7**). Taken together, our results suggest that robust detection of molecular hallmarks of TDP-43 dysfunction can occur in the absence of overt cytoplasmic TDP-43 mislocalization and aggregation in real human disease. Most significantly, these studies demonstrate that iPSNs can authentically mimic *in vivo* alterations in TDP-43 functionality in the same patient.

### Repair of nuclear pore complex injury restores TDP-43 function in sALS iPSNs

Although TDP-43 nuclear depletion and cytoplasmic aggregation has long been regarded as a pathological hallmark of ALS and related neurodegenerative diseases ^7,9,11,12,32^, the molecular mechanisms that contribute to TDP-43 pathology and in particular loss of nuclear TDP-43 function have remained understudied. We have recently demonstrated that the nuclear accumulation of CHMP7, an ESCRT-III protein involved in NPC and nuclear envelope surveillance and homeostasis ^44–48^, initiates NPC injury as defined by the reduction of specific Nups, beginning with the transmembrane Nup POM121, from sALS and C9orf72 NPCs. In turn, the collective reduction of 8 Nups compromises active nuclear import and precedes TDP-43 dysfunction in ALS iPSNs ^31^. We previously demonstrated that antisense oligonucleotide (ASO) mediated reduction of CHMP7 following the emergence of NPC injury, but prior to the detectable development of TDP-43 dysfunction, was sufficient to completely prevent the emergence of molecular hallmarks of TDP-43 loss of function in the small subset of C9orf72 and sALS iPSNs originally evaluated ^31^.

To address the extent to which a far larger population-like cohort of C9orf72 and sALS patient lines displayed nuclear accumulation of CHMP7 and reduction of POM121 compared to TDP-43 dysfunction, we performed immunostaining and confocal and AiryScan imaging for CHMP7 and POM121 respectively. Nuclear/cytoplasmic ratios of CHMP7 at day 18 of differentiation and number of POM121 spots at day 32 of differentiation were compared to qRT-PCR based analytics for our panel of TDP-43 loss of function associated mRNA processing defects at day 60 of differentiation (**Figure 7a**). In total, about 69% of lines examined displayed some level of CHMP7 nuclear accumulation and 79% displayed some level of POM121 reduction and TDP-43 dysfunction (**Figure 7b**). Focusing on the C9orf72 and sALS lines, 87% displayed CHMP7 nuclear accumulation and 100% displayed POM121 reduction and TDP-43 dysfunction (**Figure 7c**). Thus, given large size of our iPSN sample population, we conclude that alterations to the NPC are a uniform pathophysiological event in all C9orf72 and sALS patients. The observation that a small number of patient lines did not have evidence of CHMP7 nuclear accumulation (**Figure 7**) suggests that NPC injury and TDP-43 dysfunction can occur independent, albeit rarely, of alterations in CHMP7/ESCRT-III nuclear surveillance. Interestingly, nuclear accumulation of CHMP7 and POM121 reduction moderately correlated with the severity of TDP-43 dysfunction (**Supplemental Figure 8a-c**). However, when control iPSNs were not considered in the analysis, the extent of POM121 reduction no longer correlated with the magnitude of TDP-43 dysfunction in ALS iPSNs (**Supplemental Figure 8d**). Nuclear accumulation of CHMP7 and the severity of POM121 reduction appear to correlate in both analyses (**Supplemental Figure 8e-f**).

**Figure 7:**
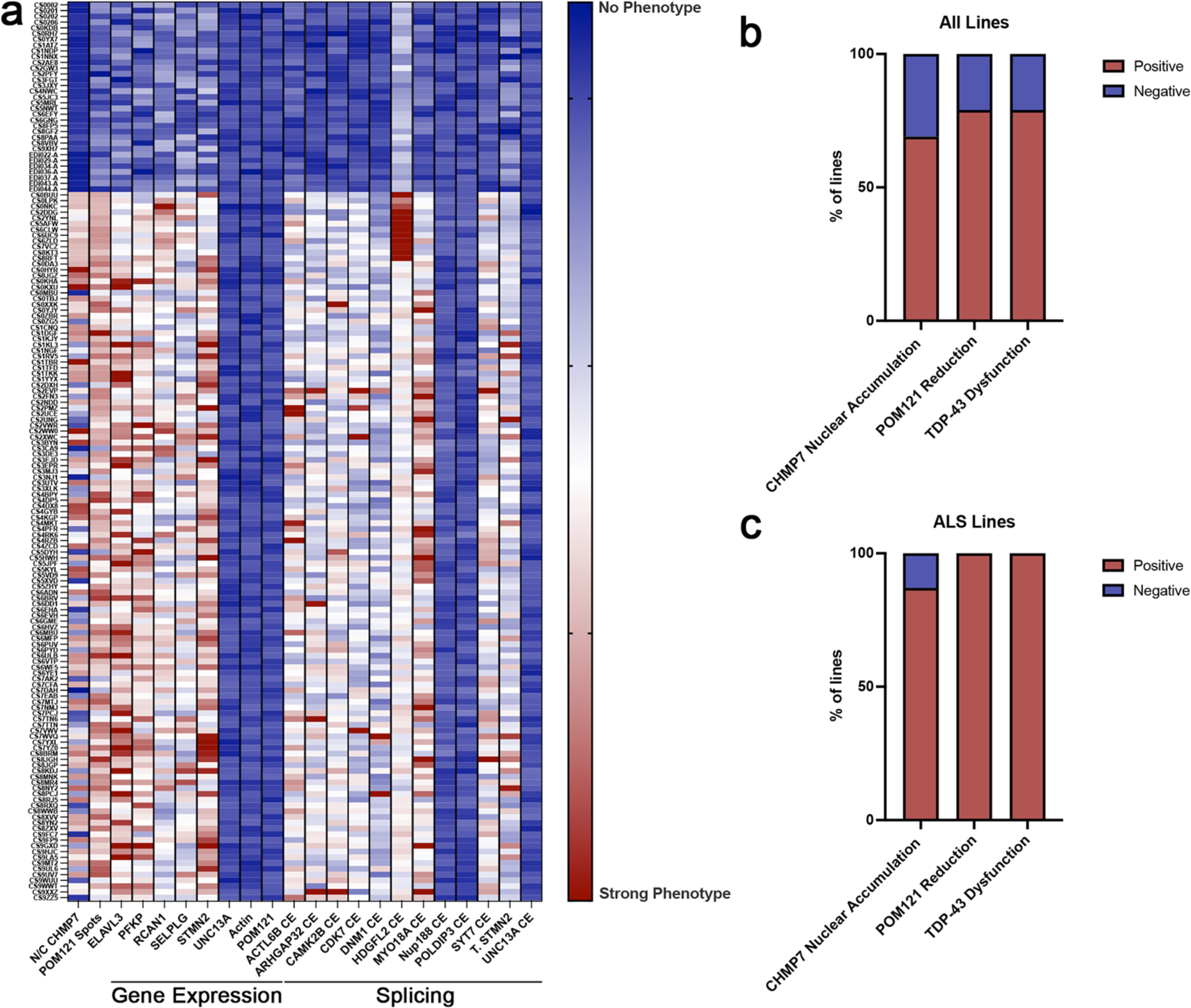
Molecular signatures of TDP-43 dysfunction correspond to hallmarks of NPC disruption in the majority of C9orf72 ALS/FTD and sALS patient iPSNs. (**a**) Heat maps of nuclear/cytoplasmic distribution of CHMP7 (obtained from confocal imaging at day 18 of differentiation), reduction of POM121 spots from the nucleus (obtained from AiryScan imaging at day 32 of differentiation), and molecular signatures of TDP-43 loss of function (obtained from qRT-PCR at day 60 of differentiation) in control, C9orf72, and sALS iPSNs. (**b**) Percentage of control, C9orf72, and sALS iPSN lines displaying CHMP7 nuclear accumulation, reduction of POM121, and TDP-43 dysfunction. (**c**) Percentage of C9orf72 and sALS iPSN lines displaying CHMP7 nuclear accumulation, reduction of POM121, and TDP-43 dysfunction. n = 33 control, 12 C9orf72, and 111 sALS iPSC lines. For each line and time point, boxes within heat map represent the average value across 3 independent differentiations.

We next treated a subset of these iPSC lines with CHMP7 ASOs ^31^ beginning at day 60 of differentiation *after* the emergence of molecular hallmarks of TDP-43 loss of function. Following 3 weeks of treatment with CHMP7 ASOs yielding sufficient reduction in CHMP7 protein (^31^ and **Supplemental Figure 9**), we observed a near complete restoration of TDP-43 function across all mRNA targets and lines evaluated (**Supplemental Figure 9**). Together with our previously published results, this suggests that reduction of CHMP7 protein can both *prevent* ^31^ and *reverse* TDP-43 dysfunction in ALS iPSNs.

Having observed that 100% of C9orf72 and sALS patient lines display POM121 reduction and molecular hallmarks of TDP-43 loss of function, we next asked whether NPC injury alone was sufficient to initiate TDP-43 dysfunction in iPSNs. To test this, we initiated NPC injury by using Trim Away ^49^ to rapidly degrade POM121 protein from otherwise control iPSNs as we have previously done ^43^. Importantly, the reduction of POM121 via Trim Away impacts NPC composition and function but is not overtly inherently toxic in the time frame of our experiments ^43^. Using qRT-PCR, we observed a significant dysregulation of TDP-43 mRNA targets across our panel of gene expression and mRNA splicing changes associated with TDP-43 function (**Supplemental Figure 10**) in control iPSNs following 7 days of POM121 reduction. Thus, these data suggest that POM121 mediated NPC injury is sufficient to initiate TDP-43 dysfunction, similar to the profile of that observed in sALS and C9orf72 iPSNs.

Given our previous report that replenishment of POM121 via overexpression following initial POM121 reduction was sufficient to re-establish NPC composition and active nuclear import in C9orf72 iPSNs ^43^, we hypothesized that POM121 overexpression may alleviate TDP-43 dysfunction which we have now established (**Supplemental Figure 10**) is one of likely many consequences of NPC injury. Therefore, we overexpressed POM121 in sALS iPSNs at day 46 of differentiation, following initial reduction and the beginning of detectable TDP-43 dysfunction. Two weeks later, consistent with CHMP7 nuclear accumulation occurring upstream of NPC injury in ALS, we did not observe a reversal of CHMP7 nuclear accumulation (**Supplemental Figure 11**). In contrast, we did detect a near complete restoration of TDP-43 function in sALS iPSNs as evaluated by our multi-target qRT-PCR panel (**Figure 8**). The restoration of TDP-43 function corresponded with a repair of active nuclear import (**Supplemental Figure 12**) as examined by the localization of a previously described and utilized S-tdTomato NCT reporter ^30,43,50^ as well as an increase in nuclear and concomitant decrease in cytoplasmic TDP-43 immunoreactivity (**Supplemental Figure 13**). Together, these results suggest that NPC injury, downstream of CHMP7 nuclear accumulation, can on its own directly impact TDP-43 function and repair of NPC composition and function itself may be a viable therapeutic strategy for alleviating TDP-43 dysfunction in sALS.

**Figure 8:**
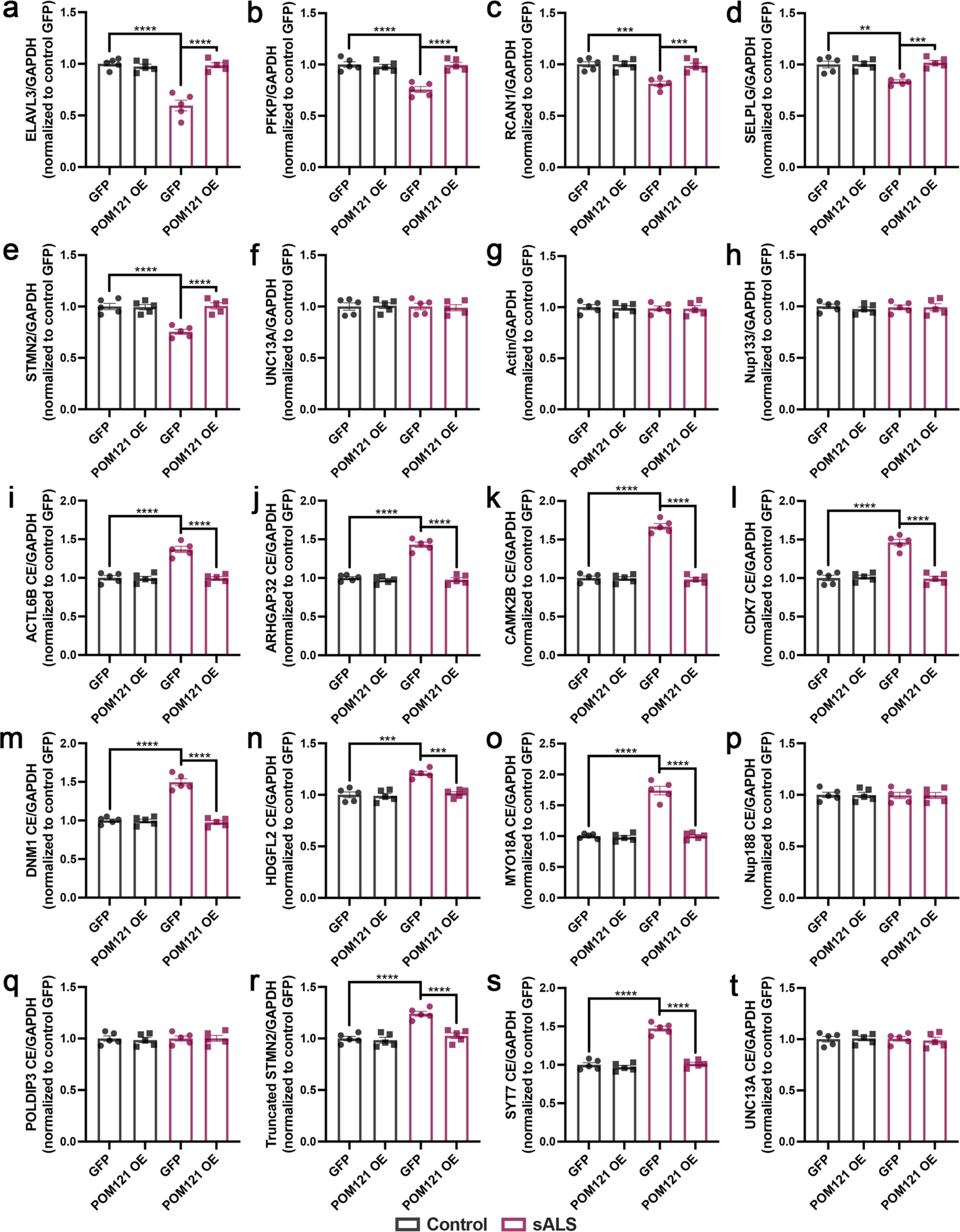
Overexpression of the transmembrane Nup POM121 is sufficient to restore TDP-43 function in sALS iPSNs. (**a-t**) qRT-PCR for *ELAVL3* (**a**), *PFKP* (**b**), *RCAN1* (**c**), and *SELPLG* (**d**), *STMN2* (**e**), *UNC13A* (**f**), *ACTIN* (**g**), *POM121* (**h**), *ACTL6B* cryptic exon containing (**i**), *ARHGAP32* cryptic exon containing (**j**), *CAMK2B* cryptic exon containing (**k**), *CDK7* cryptic exon containing (**l**), *DNM1* cryptic exon containing (**m**), *HDGFL2* cryptic exon containing (**n**), *MYO18A* cryptic exon containing (**o**), *NUP188* cryptic exon containing (**p**), *POLDIP3* cryptic exon containing (**q**), *SYT7* cryptic exon containing (**r**), truncated *STMN2* (**s**), and *UNC13A* cryptic exon containing (**t**) mRNA in control and sALS iPSNs at day 67 of differentiation following 3 weeks of GFP or POM121 overexpression. POM121 overexpression was initiated at day 46 of differentiation following the detectable initiation of TDP-43 dysfunction in sALS iPSNs. GAPDH was used as the reference gene for normalization. *ACTIN* and *POM121* were used as negative control mRNAs not known to be regulated by TDP-43. n = 5 control and 5 sALS iPSC lines. Two-way ANOVA with Tukey’s multiple comparison test was used to calculate statistical significance. ** p < 0.01, *** p < 0.001, **** p < 0.0001.

## Discussion

The loss of nuclear TDP-43 function is widely regarded as an early and significant contributor to disease pathogenesis in ALS and related neurodegenerative diseases ^18–27^. However, authentic preclinical models that recapitulate this event have been challenging to define. Recent advances in the availability of sALS patient iPSC lines ^29^ and iPSN differentiation methodology provide an unparalleled opportunity to begin to study ALS pathophysiology in authentic human neuronal models of sporadic disease. However, examining pathophysiologic events in inherently heterogenous models of sALS has proven to be a daunting task. Here we examine 6 gene expression changes and 12 cryptic exon splicing events previously linked to TDP-43 depletion ^19,20,25^ in >150 iPSC lines at 3 time points in iPSN differentiation in total generating ∼30,000 qRT-PCR based data points. By studying such a large population of patient derived iPSC lines, we provide a far clearer view of both the real consistency of pathophysiological events (e.g. CHMP7 nuclear accumulation, NPC injury, overall TDP-43 dysfunction), but also the great variability that accompanies sporadic disease. In doing so, this study revealed a significant time dependent emergence of molecular signatures of TDP-43 dysfunction (**Figures 1-4, Supplemental Figures 1-2**) in authentic sALS patient iPSNs. However, not all gene expression or splicing alterations previously linked to TDP-43 depletion were observed at the time points evaluated (**Figures 1-4, Supplemental Figures 1-2**). Strikingly, we observed an extensive variability in the magnitude of gene expression and cryptic exon containing mRNA detection both across mRNAs examined in individual patient iPSC lines and amongst patient iPSC lines for individual mRNAs (**Figures 1-4, Supplemental Figures 1-2**). Although the mechanisms underlying this variability remain unclear at this time, it is possible that SNPs (e.g. similar to that reported for UNC13A ^20,22^) or co-regulation of mRNA targets by multiple RNA binding proteins (as has been previously demonstrated for TDP-43 and FMRP in *Drosophila* models ^51,52^) may be contributing factors. Nonetheless, this variability holds implications for future sALS focused clinical trials either targeting individual mRNA targets of TDP-43 themselves ^19,23,27,28^ or those seeking to utilize TDP-43 loss of function associated “cryptic” peptides or RNAs as biomarkers or readouts of therapeutic efficacy ^21,25^.

Some might “argue” that iPSC models reflect only immature neuronal biology. Therefore, the comparison of a biological pathway in iPSN to actual human brain from the same patient becomes quite informative. Critically, the “TDP-43 dysfunction” signatures we detected are similar in postmortem motor cortex and spinal cord tissues compared to iPSNs from the same patients (**Figure 6**) demonstrating the utility of our iPSN model in recapitulating events that occur in real human disease.

Nuclear depletion of TDP-43 is often associated with cytoplasmic mislocalization and/or aggregation ^7,12,33^. Thus, it stands to reason that nuclear loss of function would be dependent upon at least some detectable level of nuclear depletion. However, we clearly demonstrate that TDP-43 dysfunction can be robustly detected in bulk cellular and tissue lysates (**Figures 1-6, Supplemental Figures 1-2**) in the absence of overt histological nuclear clearance in the majority of cells even in postmortem tissues (**Figure 6, Supplemental Figure 3**). Moreover, consistent with prior reports ^14–16^, cytoplasmic TDP-43 aggregates were rare in end-stage disease tissues (**Figure 6**). Thus, these data suggest that histologically, TDP-43 may remain nuclear and yet not appropriately engage in its typical nuclear functions. Although the mechanisms that may underlie this potential phenomenon are not yet understood, these data underscore the notion that loss of nuclear function does not necessarily require a complete loss of nuclear compartmentalization or overt aggregation.

Although TDP-43 pathology has been documented in ALS, and more recently related neurodegenerative disease, for nearly two decades ^7–16^ and loss of nuclear TDP-43 function is considered to be a significant contributor to disease pathophysiology ^9,18,32,53–56^, the mechanisms that contribute to these events are poorly understood. We have previously demonstrated that the pathologic nuclear accumulation of the ESCRT-III nuclear surveillance protein CHMP7 is a significant contributor to NPC injury events and subsequent TDP-43 dysfunction in sALS and C9orf72 ALS/FTD ^31^. In this current study, we now provide evidence that NPC injury itself can directly contribute to altered TDP-43 functionality in sALS iPSNs (**Supplemental Figure 10**). Given that the NPC directly governs NCT and the NPC and its Nup constituents directly and indirectly control numerous cellular processes via regulation of genome organization and gene expression ^57–61^, injury to the NPC or Nups is likely to have wide-ranging implications for cellular function. Notably, at this time, it remains unclear how and what aspect of NPC injury directly impacts TDP-43 function. Nevertheless, overexpression or artificial replacement of the transmembrane Nup POM121, a central player in NPC injury cascades ^43^, resulted in a near complete restoration of TDP-43 function in sALS iPSNs (**Figure 8**). Thus, in contrast to recent strategies that aim to restore the expression of single TDP-43 targets ^19,23,27,28^, these data highlight the potential for directly targeting the NPC as a therapeutic strategy for restoring the expression of *multiple* mRNA targets implicated in TDP-43 loss of function events.

In summary, here we provide evidence for extensive variability in detected molecular signatures of TDP-43 dysfunction within and amongst individual sALS patient iPSC lines that recapitulates signatures from postmortem CNS tissues. This data set demonstrates the utility of iPSN models for studying pathophysiological events in sALS and for preclinical therapeutic discovery and testing and has important implications for future clinical trial design. Importantly, our characterization of TDP-43 dysfunction in >150 iPSC lines represents an essential resource for the community for selection of lines for future studies.

## Methods

### iPSC Differentiation

All iPSC lines used in this study detailed in **Supplemental File 2** were obtained from the Answer ALS ^29^ repository at Cedars Sinai. iPSCs were maintained in mTeSR Plus media as recently described ^30^. Information on iPSC lines displaying the most robust overall TDP-43 dysfunction score as well as those with the highest magnitude of change for each mRNA species examined can be found in **Supplemental File 3**. Mixed spinal neuron cultures were generated using a modified direct induced motor neuron (diMNs) protocol that has recently been described in detail ^30^. Cortical neuron cultures were generated as recently detailed ^38^ with no protocol modifications. To generate relatively pure populations of “cortical-like” neurons and lower motor neurons, we utilized PiggyBac integration technology to express PB-tet-NGN2 (Addgene 172115; “cortical-like” neurons) and PB-tet-hNIL (Addgene 172113; lower motor neurons) as recently described ^62,63^ with no protocol modifications. All iPSC and iPSN cultures were maintained at 37°C with 5% CO2 and routinely tested negative for mycoplasma.

### CHMP7 ASO treatment in iPSNs

Non-targeting scrambled control (676630): CCTATAGGACTATCCAGGAA and CHMP7 ASO (1508917): TGTTACCCTCAGATACCGCC were generously provided by Ionis Pharmaceuticals and have been previously described ^31^. On day 60 of differentiation, ASOs were added to iPSN media to a final concentration of 5 μM. Every 3-4 days, media was exchanged and ASO was replaced until the experimental time point indicated in figure legends.

### RNA isolation, cDNA synthesis, and qRT-PCR

iPSN Preparation: On the day of isolation, 1 mL of Trizol was added to each well of iPSNs in a 6 well plate. Following a 5 minute incubation at room temperature, Trizol/lysed cell solutions were transferred to an Eppendorf tube. Postmortem Human Tissue Preparation: 25 mg frozen tissue sections were homogenized in 1 mL Trizol with a dounce homogenizer. Trizol/tissue lysates were transferred to an Eppendorf tube. For iPSNs and postmortem human tissues, Trizol based RNA isolation proceeded in accordance with manufacturer protocol (Invitrogen). cDNA synthesis was carried out using the High Capacity cDNA Reverse Transcription Kit (Thermo Fisher Scientific). 1 μg of RNA was used for each reaction. All qRT-qPCR reactions were conducted using SYBR Green Master Mix or TaqMan Gene Expression Master Mix (Thermo Fisher) and an Applied Biosystems QuantStudio 3 (Applied Biosystems). Previously described primer sets (see **Supplemental File 4** for sequences) ^22,23,25,64^ were used to detect truncated STMN2 and cryptic exon containing mRNA transcripts. TaqMan Gene Expression Assays (see **Supplemental File 4** for probe information) were used to detect mRNA targets. GAPDH was used for normalization of gene expression.

### Human tissue immunostaining and imaging

Paraffin embedded sALS patient postmortem motor and occipital cortex tissue sections (see **Supplemental File 5**) were deparaffinized, immunostained, and imaged as previously described ^30,31^. Primary antibodies used were rabbit anti-TDP-43 (ProteinTech 10782-2-AP) and guinea pig anti-Map2 (Synaptic Systems 188004). Goat anti-rabbit Alexa 488 and goat anti-guinea pig Alexa 568 (Thermo Fisher Scientific) were used as secondary antibodies. Tissue sections were imaged with a 20X objective on a Zeiss Axioimager Z2 fluorescent microscope housing an apotome2 module. All images were acquired with identical exposure times. Categorical analysis was carried about by manual observation and counting of cells displaying strong nuclear TDP-43 immunoreactivity, strong nuclear with cytoplasmic TDP-43 immunoreactivity, weak nuclear TDP-43 immunoreactivity, nuclei completely devoid of TDP-43 immunoreactivity, and cells with visible cytoplasmic TDP-43 aggregates. Nuclear/cytoplasmic ratios of TDP-43 were quantified as previously described ^31^ whereby integrated density was measured within a nuclear ROI and cytoplasmic ROI within the cell body. Background mean intensity values were subtracted prior to calculating nuclear/cytoplasmic ratios.

### Plasmid expression and Trim Away

Plasmids were expressed in iPSNs via suspension based nucleofection with the Lonza 4D nucleofection system as previously described in detail ^30,31,43^ at time points indicated in figure legends. GFP, POM121 GFP, and S-tdTomato plasmids have been previously described ^30,43^. To enrich for iPSNs expressing GFP or POM121 GFP (Origene), cultures were exposed to neomycin based selection every third day for the duration of the experiment following nucleofection. Trim Away ^49^ based degradation of POM121 in iPSNs was carried out as previously described ^43^ at time point indicated in figure legends.

### Immunostaining and confocal imaging of iPSNs

Five days prior to fixation, iPSNs were dissociated with accutase and re-plated in 24-well glass bottom plates as recently described ^30^. Immunostaining and confocal imaging were carried out as previously described ^30,31^. For all experiments, POM121 GFP overexpression was visualized by native GFP expression. For TDP-43 detection, primary antibodies used were rabbit anti-TDP-43 (ProteinTech 10782-2-AP) and guinea pig anti-Map2 (Synaptic Systems 188004). Goat anti-rabbit Alexa 647 and goat anti-guinea pig Alexa 568 (Thermo Fisher Scientific) were used as secondary antibodies. Following addition of mounting media and coverslips, iPSNs were imaged with a 63X objective on a Zeiss LSM 980 confocal microscope. For immunostaining of CHMP7, mouse anti-CHMP7 (Santa Cruz sc-271805) and guinea pig anti-Map2 (Synaptic Systems 188004) were used as primary antibodies and Goat anti-rabbit Alexa 647 and goat anti-guinea pig Alexa 568 (Thermo Fisher Scientific) were used as secondary antibodies. For imaging of the S-tdTomato NCT reporter, iPSNs were fixed in 4% PFA, washed with 1X PBS (3X 10 mins) and incubated with Hoescht (1:1000 in 1X PBS) for 10 minutes prior to additional washing and immediate imaging with a 63X objective on a Zeiss LSM 980 confocal microscope as recently described ^30^. All images within each data set (CHMP7, TDP-43, S-tdTomato) were acquired using identical imaging parameters including frame scanning speed, laser power, and gain.

### Statistical Analysis

All image analysis was blinded. Statistical analyses were performed using GraphPad Prism version 10 (GraphPad). As previously described, for imaging-based experiments in iPSNs, the average values for all cells analyzed per iPSC was considered n = 1 ^30,31,43^. The total number of cells evaluated per experiment is reported in the figure legends. As appropriate for the experimental design, Student’s t-test, One-way or Two-way ANOVA with Tukey’s multiple comparison test was used as described in figure legends. * p < 0.05, ** p < 0.01, *** p < 0.001, **** p < 0.0001. Violin plots are used to display the full spread and variability of data within imaging analysis of iPSNs. The center dotted line indicates the median value. Two additional dotted lines indicate the 25^th^ and 75^th^ percentiles. Individual data points represent the average value of all cells analyzed per iPSC line. Bar graphs with individual data points representing each iPSC line, histograms or stacked bars representing percentage of cells for human tissue analyses, or heat maps and line graphs are used to display data obtained from all other assays and as detailed in figure legends.

## Acknowledgements

We thank the ALS patients and their families whose contributions made this research possible. iPSC lines were acquired through the Answer ALS repository at Cedars Sinai (https://csbiomfg.com/cellcollection). Postmortem human tissues were obtained from Jonathan Glass and the Goizueta Alzheimer’s Disease Center at Emory University (P30 AG066511), Cindy Ly and Timothy Miller at Washington University, and the Johns Hopkins postmortem tissue bank (Jeffrey Rothstein). This work was supported by The Robert Packard Center for ALS Research (ANC), NIH NINDS/NIA R00 NS123242 (ANC), NIH NINDS R01 NS132836 (ANC), Answer ALS (JDR), NIH NIA R01 RF1AG062171 (JDR), Chan Zuckerberg Foundation (JDR), NIH NINDS 2P01NS084974, R01 NS122236 (JDR) R35 NS132179 (JDR), ALS Association (JDR), Muscular Dystrophy Association (JDR), Virginia Gentlemen Foundation (JDR), US Dept of Defense HT94252310136 (JDR), F Prime (JDR).

## Author Contributions

Conceived and designed the experiments: ANC. Performed the experiments: ANC and CW. Analyzed the data: ANC and JDR. Contributed reagents and materials: ANC and JDR. Wrote the manuscript: ANC. Edited the manuscript: ANC and JDR. Reviewed the final version of the manuscript: ANC, CW, and JDR.

## Figures and Legends

**Supplementary Figure 1:**
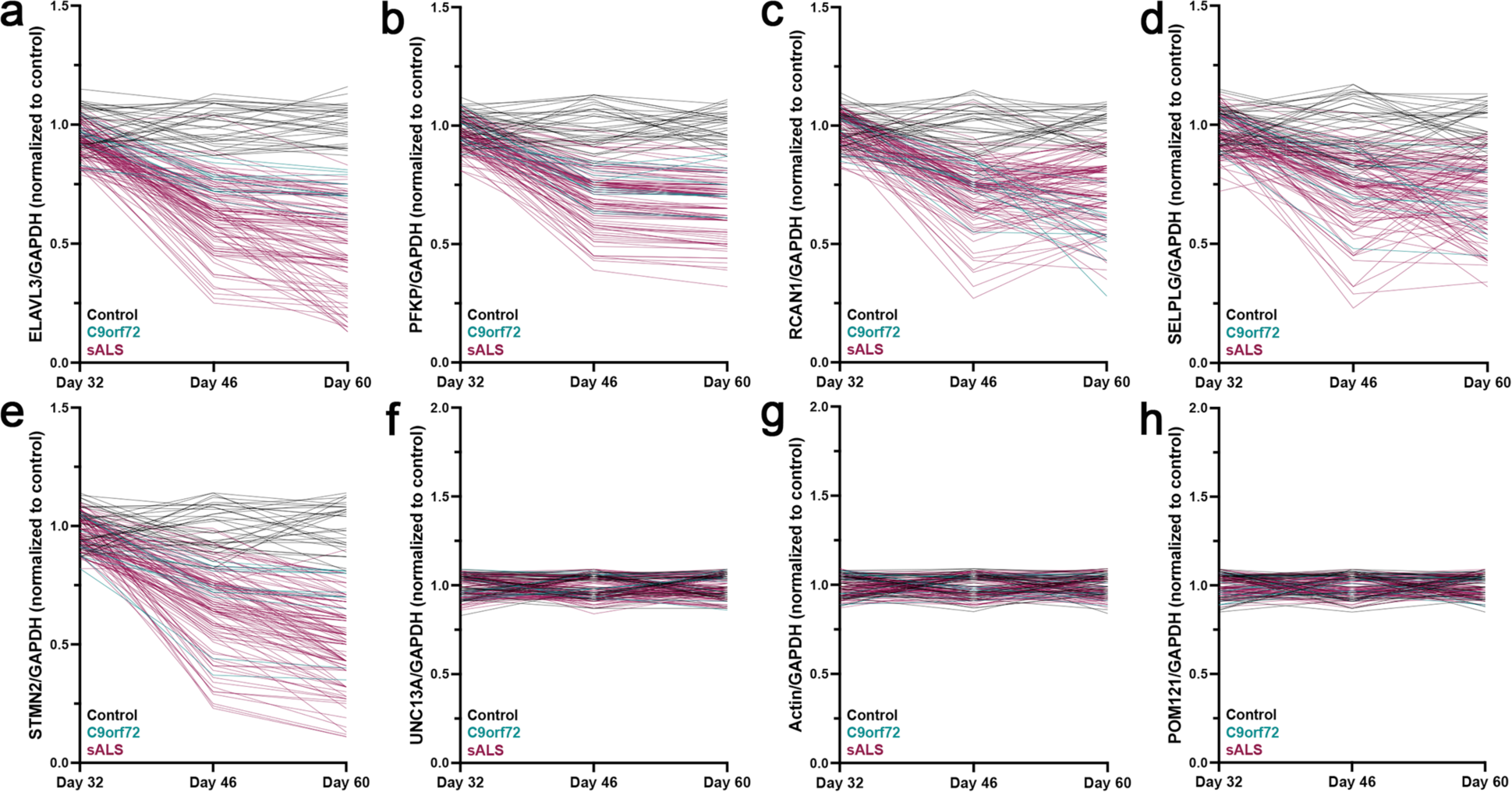
Time dependent and patient specific profiles of TDP-43 loss of function associated changes in target gene expression in C9orf72 ALS/FTD and sALS iPSNs. **(a-h)** Line graphs from qRT-PCR for *ELAVL3* **(a)**, *PFKP* **(b)**, *RCAN1* **(c)**, and *SELPLG* **(d)**, *STMN2* **(e)**, *UNC13A* **(f)**, *ACTIN* **(g)**, and *POM121* **(h)** mRNA in control, C9orf72, and sALS iPSNs at day 32, 46, and 60 of differentiation. *ACTIN* and *POM121* represent negative control mRNAs not known to be regulated by TDP-43. n = 33 control, 12 C9orf72, and 111 sALS iPSC lines. For each line and time point, data points represent the average value across 3 independent differentiations.

**Supplementary Figure 2:**
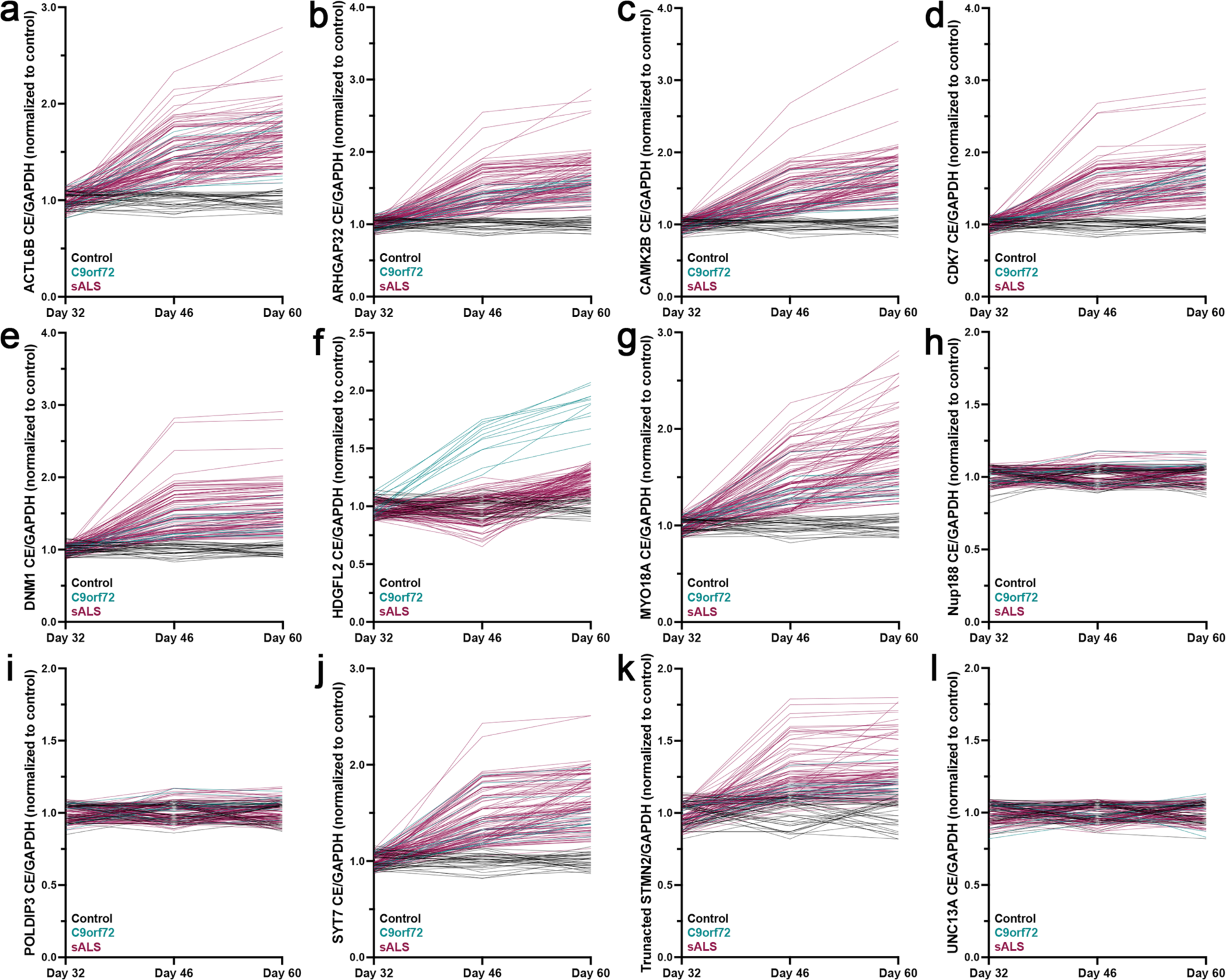
Time dependent and patient specific profiles of TDP-43 loss of function associated changes in target mRNA splicing in C9orf72 ALS/FTD and sALS iPSNs. Line graphs from qRT-PCR for *ACTL6B* cryptic exon containing. **(a)**, *ARHGAP32* cryptic exon containing **(b)**, *CAMK2B* cryptic exon containing **(c)**, *CDK7* cryptic exon containing **(d)**, *DNM1* cryptic exon containing **(e)**, *HDGFL2* cryptic exon containing **(f)**, *MYO18A* cryptic exon containing **(g)**, *NUP188* cryptic exon containing **(h)**, *POLDIP3* cryptic exon containing **(i)**, *SYT7* cryptic exon containing **(j)**, truncated *STMN2* **(k)**, and *UNC13A* cryptic exon containing **(l)** mRNA in control, C9orf72, and sALS iPSNs at day 32, 46, and 60 of differentiation. n = 33 control, 12 C9orf72, and 111 sALS iPSC lines. For each line and time point, data points represent the average value across 3 independent differentiations.

**Supplementary Figure 3:**
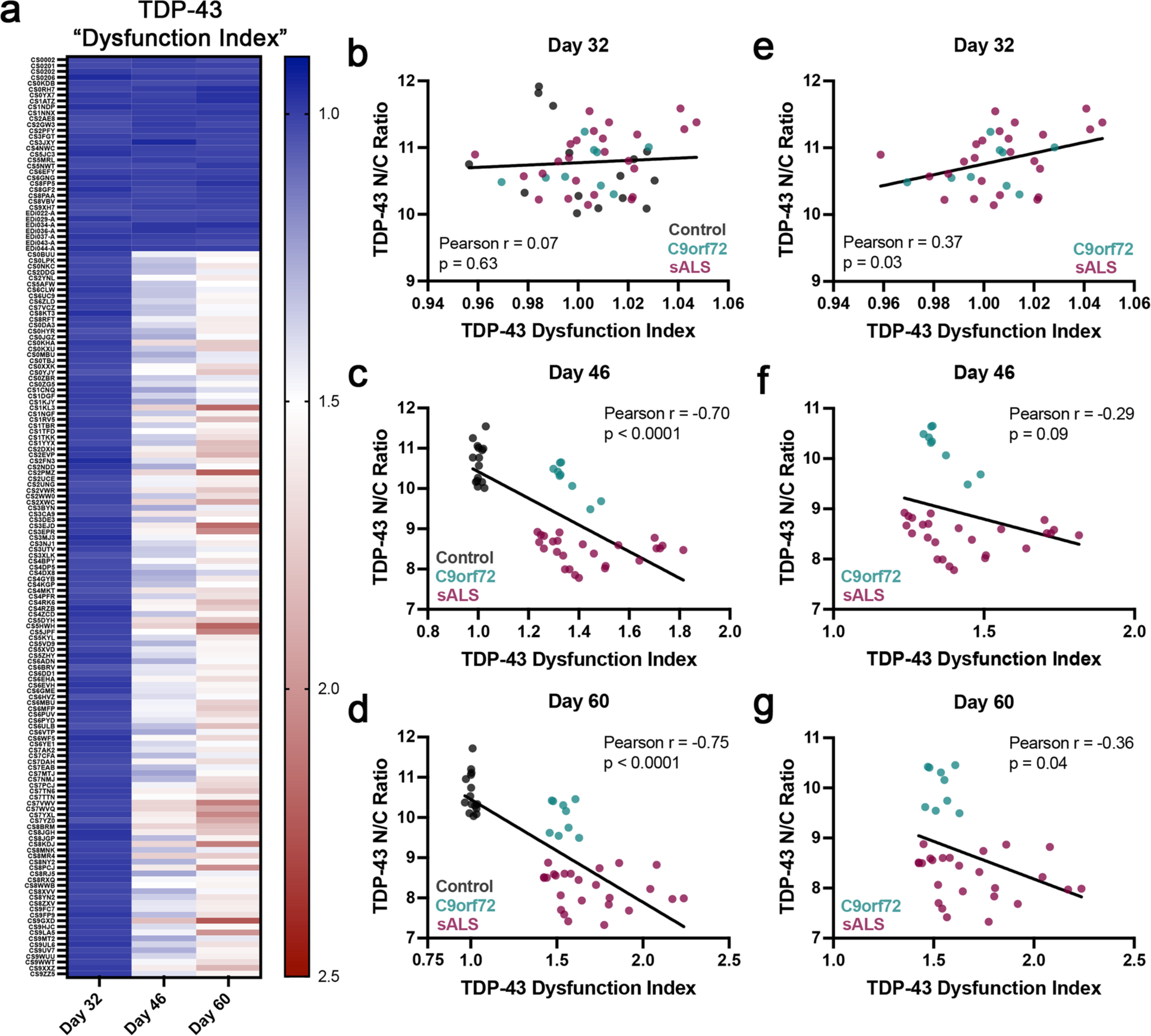
Time dependent molecular signatures of TDP-43 dysfunction correlate with subtle decreased in nuclear TDP-43 localization. **(a)** Heat map depicting overall magnitude of TDP-43 dysfunction combined from qRT-PCR for a panel of 20 TDP-43 loss of function associated alterations in target gene expression and mRNA splicing in control, C9orf72, and sALS iPSNs at day 32, 46, and 60 of differentiation. n = 33 control, 12 C9orf72, and 111 sALS iPSC lines. For each line and time point, boxes represent a composite “score” obtained from the average value across 3 independent differentiations for each of the 20 target genes evaluated. **(b-d)** Nuclear/cytoplasmic distribution of TDP-43 (obtained from immunostaining and confocal imaging) compared to overall magnitude of TDP-43 dysfunction in control, C9orf72, and sALS iPSNs at day 32 **(b)**, day 46 **(c)**, and day 60 **(d)** of differentiation. **(e-g)** Nuclear/cytoplasmic distribution of TDP-43 (obtained from immunostaining and confocal imaging) compared to overall magnitude of TDP-43 dysfunction in C9orf72 and sALS iPSNs at day 32 **(e)**, day 46 **(f)**, and day 60 **(g)** of differentiation.

**Supplementary Figure 4:**
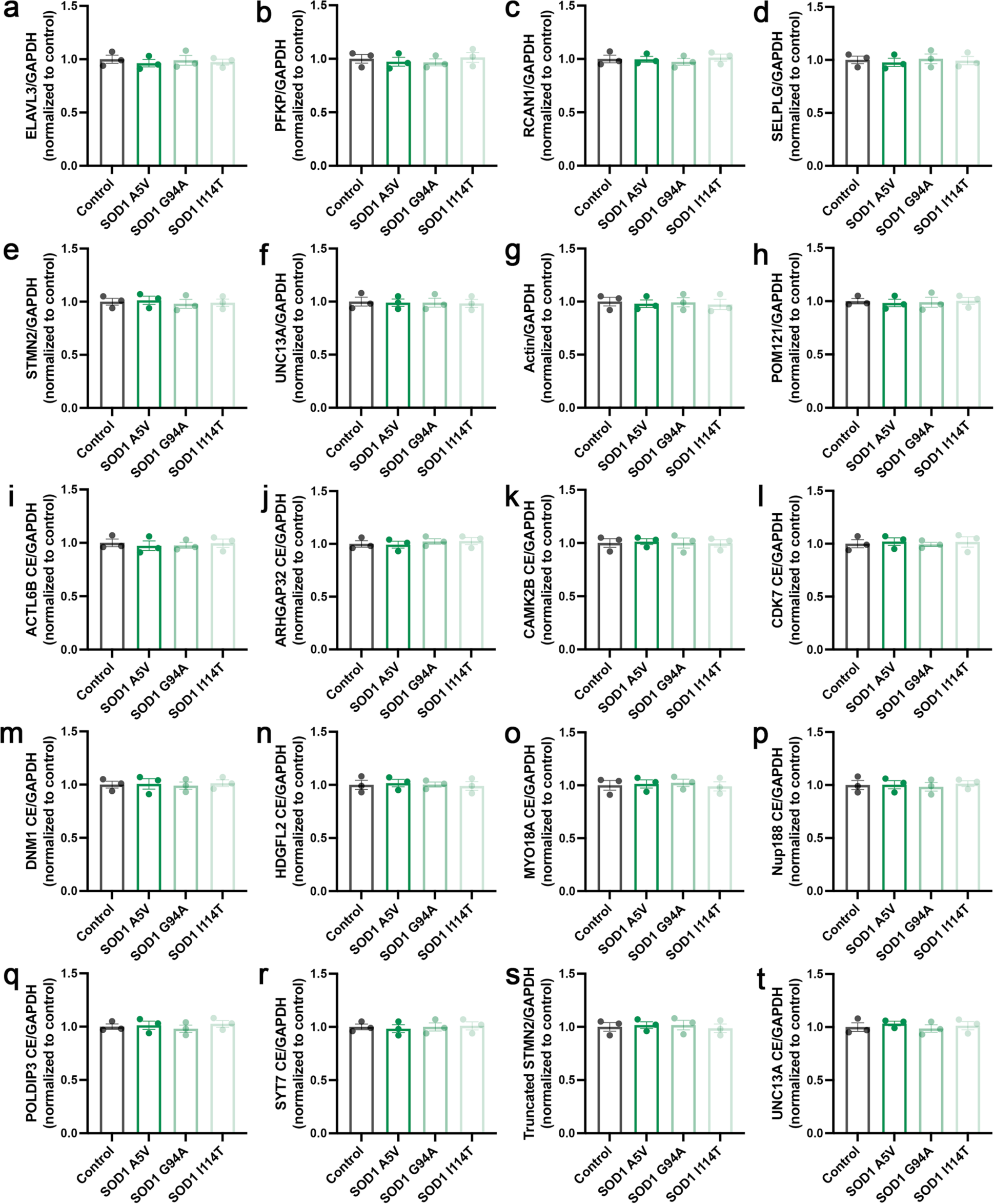
TDP-43 loss of function associated changes in target gene expression and mRNA splicing are not observed in SOD1 patient iPSNs. **(a-t)** qRT-PCR for *ELAVL3* **(a)**, *PFKP* **(b)**, *RCAN1* **(c)**, and *SELPLG* **(d)**, *STMN2* **(e)**, *UNC13A* **(f)**, *ACTIN* **(g)**, *POM121* **(h)**, *ACTL6B* cryptic exon containing **(i)**, *ARHGAP32* cryptic exon containing **(j)**, *CAMK2B* cryptic exon containing **(k)**, *CDK7* cryptic exon containing **(l)**, *DNM1* cryptic exon containing **(m)**, *HDGFL2* cryptic exon containing **(n)**, *MYO18A* cryptic exon containing **(o)**, *NUP188* cryptic exon containing **(p)**, *POLDIP3* cryptic exon containing **(q)**, *SYT7* cryptic exon containing **(r)**, truncated *STMN2* **(s)**, and *UNC13A* cryptic exon containing **(t)** mRNA in control and SOD1 mutant iPSNs at day 60 of differentiation. GAPDH was used as the reference gene for normalization. *ACTIN* and *POM121* were used as negative control mRNAs not known to be regulated by TDP-43. n = 3 control and 3 SOD1 mutant iPSC lines. Control data points represent the average value across 3 independent differentiations for each line. Each SOD1 mutation represented by individual bars. SOD1 data points represent values for each of 3 independent differentiations per mutant line.

**Supplementary Figure 5:**
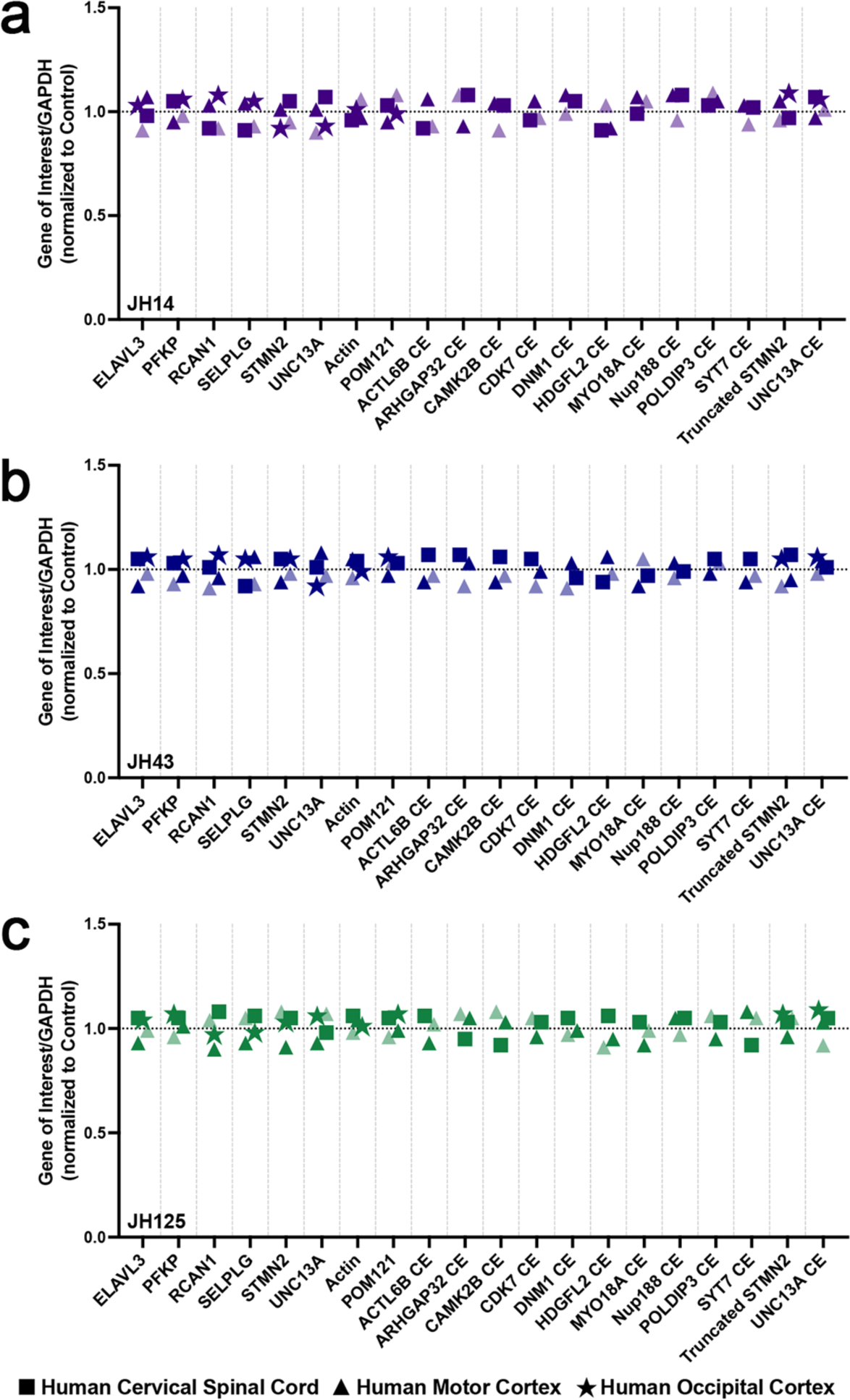
TDP-43 loss of function associated changes in target gene expression and mRNA splicing are not observed in postmortem SOD1 patient CNS tissues. **(a-c)** qRT-PCR for TDP-43 loss of function associated changes in gene expression and mRNA splicing in postmortem SOD1 patient tissues. GAPDH was used as the reference gene for normalization. *ACTIN* and *POM121* were used as negative control mRNAs not known to be regulated by TDP-43. Deidentified patient code as indicated in lower left. Symbols represent CNS region as detailed in legend. Normalized control values represented by horizontal dashed line.

**Supplementary Figure 6:**
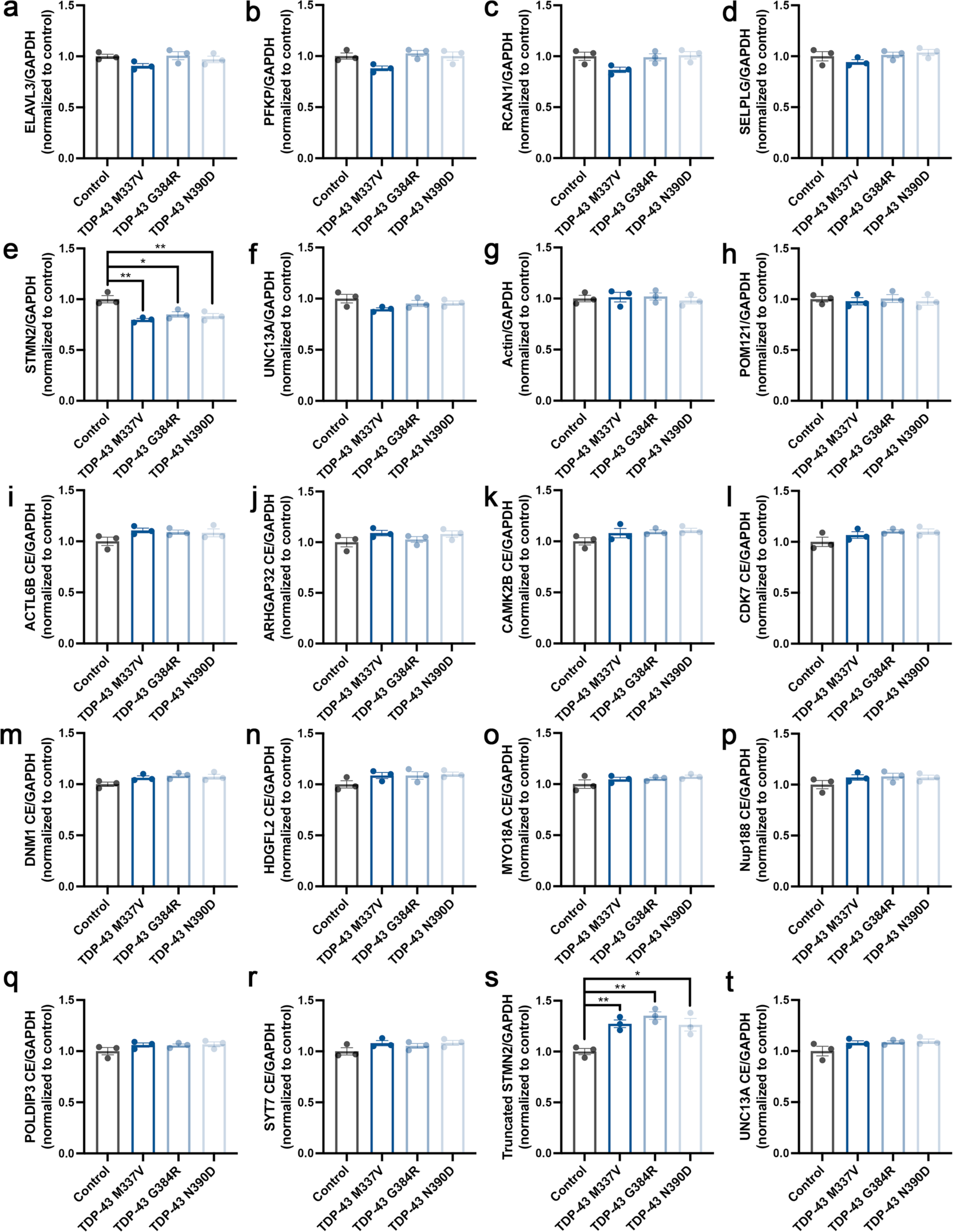
TDP-43 loss of function associated changes in STMN2 gene expression and mRNA splicing are observed in TDP-43 patient iPSNs. (a-t) qRT-PCR for *ELAVL3*. **(a)**, *PFKP* **(b)**, *RCAN1* **(c)**, and *SELPLG* **(d)**, *STMN2* **(e)**, *UNC13A* **(f)**, *ACTIN* **(g)**, *POM121* **(h)**, *ACTL6B* cryptic exon containing **(i)**, *ARHGAP32* cryptic exon containing **(j)**, *CAMK2B* cryptic exon containing **(k)**, *CDK7* cryptic exon containing **(l)**, *DNM1* cryptic exon containing **(m)**, *HDGFL2* cryptic exon containing **(n)**, *MYO18A* cryptic exon containing **(o)**, *NUP188* cryptic exon containing **(p)**, *POLDIP3* cryptic exon containing **(q)**, *SYT7* cryptic exon containing **(r)**, truncated *STMN2* **(s)**, and *UNC13A* cryptic exon containing **(t)** mRNA in control and TDP-43 mutant iPSNs at day 60 of differentiation. GAPDH was used as the reference gene for normalization. *ACTIN* and *POM121* were used as negative control mRNAs not known to be regulated by TDP-43. n = 3 control and 3 TDP-43 mutant iPSC lines. Control data points represent the average value across 3 independent differentiations for each line. Each TDP-43 mutation represented by individual bars. TDP-43 data points represent values for each of 3 independent differentiations per mutant line. One-way ANOVA with Tukey’s multiple comparison test was used to calculate statistical significance. * p < 0.05, ** p < 0.01.

**Supplementary Figure 7:**
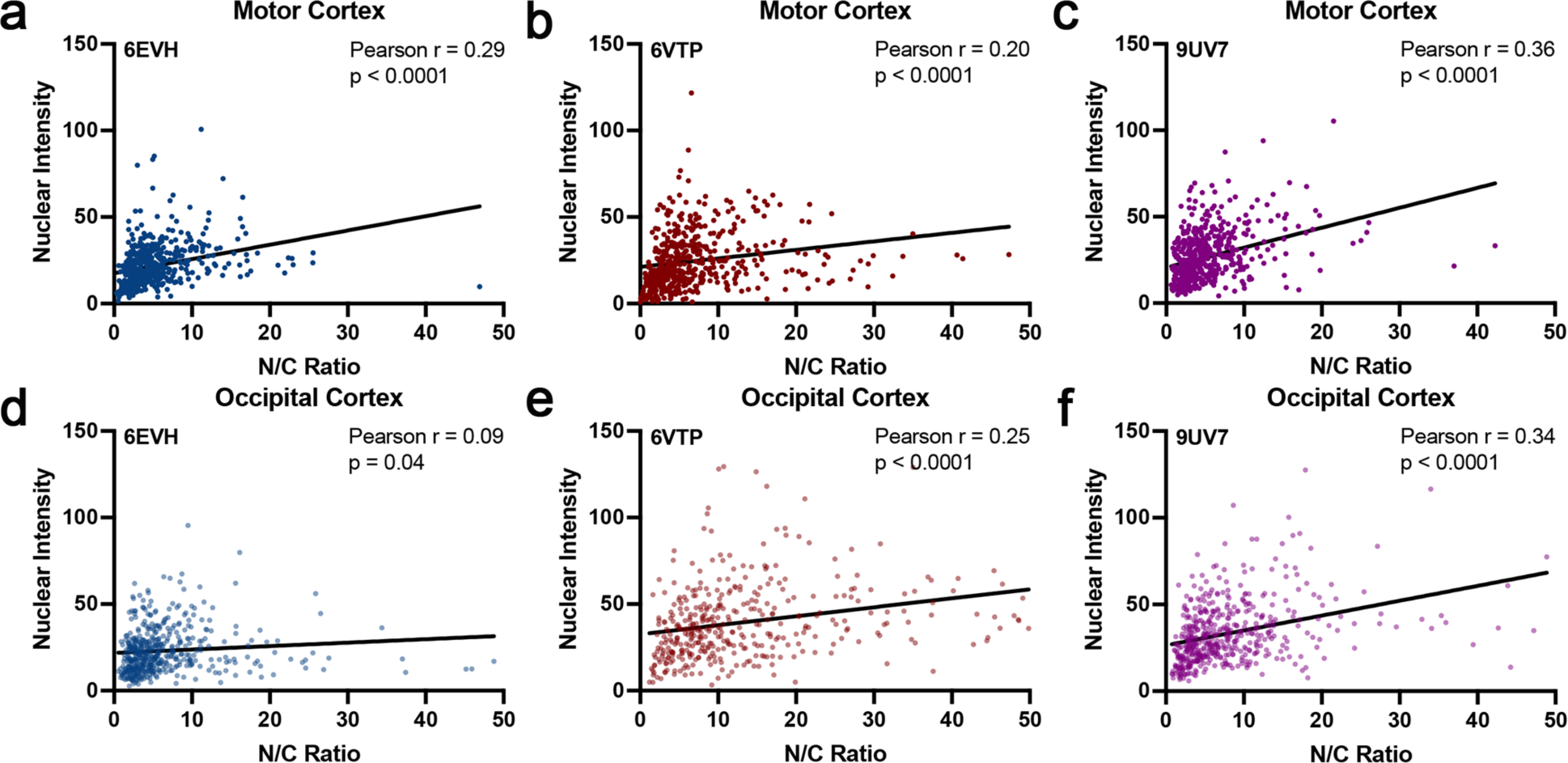
Correlation of TDP-43 nuclear/cytoplasmic distribution and nuclear intensity in postmortem sALS patient brain tissue. **(a-c)** Nuclear/cytoplasmic distribution of TDP-43 compared to nuclear intensity of TDP-43 in postmortem sALS patient motor cortex. Deidentified sALS patient code as indicated in upper left. **(d-f)** Nuclear/cytoplasmic distribution of TDP-43 compared to nuclear intensity of TDP-43 in postmortem sALS patient occipital cortex. Deidentified sALS patient code as indicated in upper left.

**Supplementary Figure 8:**
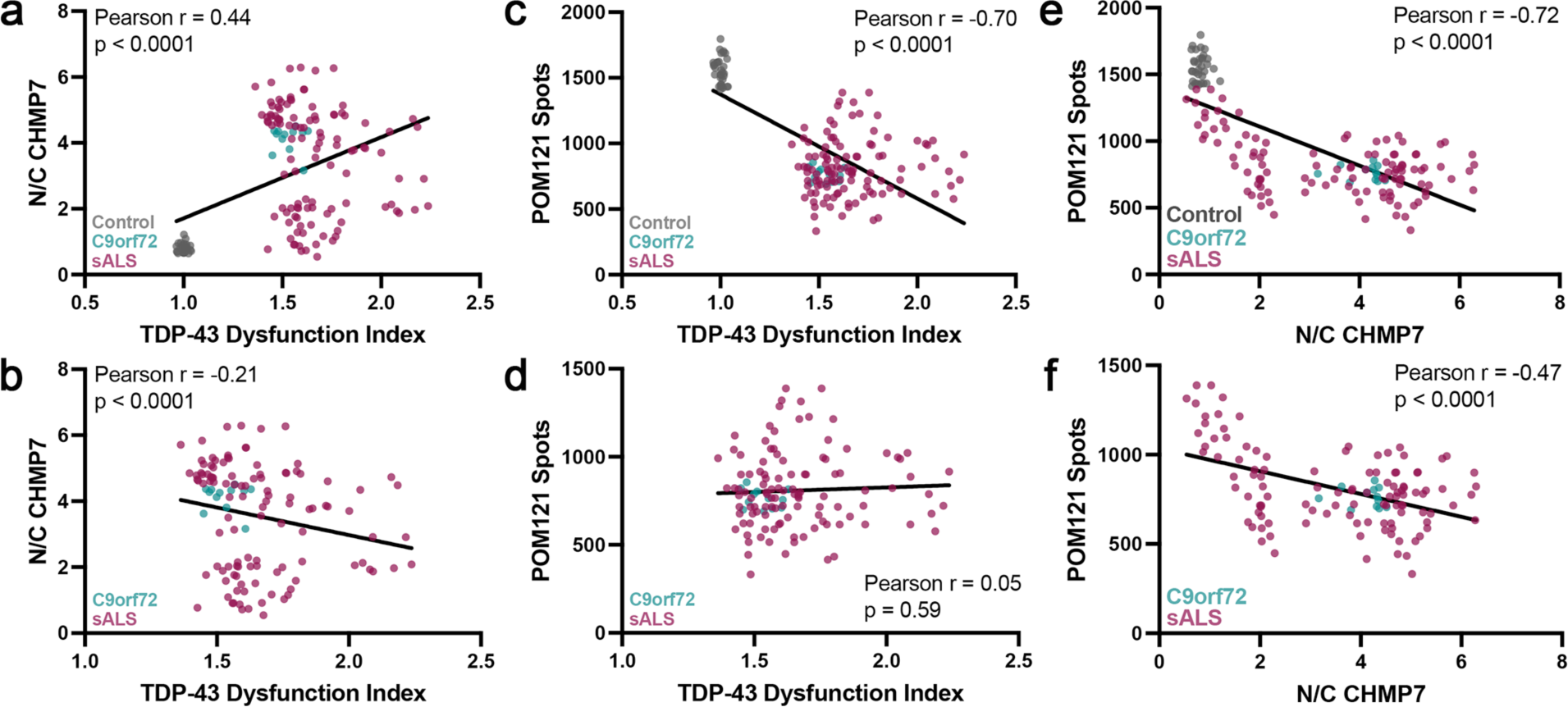
Correlation of pathological hallmarks of NPC injury events in iPSNs. **(a)** Nuclear/cytoplasmic distribution of CHMP7 (obtained from immunostaining and confocal imaging at day 18 of differentiation) compared to overall magnitude of TDP-43 dysfunction (obtained from qRT-PCR at day 60 of differentiation) in control, C9orf72, and sALS iPSNs. **(b)** Number of POM121 spots (obtained from immunostaining and AiryScan imaging at day 32 of differentiation) compared to overall magnitude of TDP-43 dysfunction (obtained from qRT-PCR at day 60 of differentiation) in control, C9orf72, and sALS iPSNs. **(c)** Number of POM121 spots (obtained from immunostaining and AiryScan imaging at day 32 of differentiation) compared to nuclear/cytoplasmic distribution of CHMP7 (obtained from immunostaining and confocal imaging at day 18 of differentiation) in control, C9orf72, and sALS iPSNs. **(d)** Nuclear/cytoplasmic distribution of CHMP7 (obtained from immunostaining and confocal imaging at day 18 of differentiation) compared to overall magnitude of TDP-43 dysfunction (obtained from qRT-PCR at day 60 of differentiation) in C9orf72 and sALS iPSNs. **(e)** Number of POM121 spots (obtained from immunostaining and AiryScan imaging at day 32 of differentiation) compared to overall magnitude of TDP-43 dysfunction (obtained from qRT-PCR at day 60 of differentiation) in C9orf72 and sALS iPSNs. **(f)** Number of POM121 spots (obtained from immunostaining and AiryScan imaging at day 32 of differentiation) compared to nuclear/cytoplasmic distribution of CHMP7 (obtained from immunostaining and confocal imaging at day 18 of differentiation) in C9orf72 and sALS iPSNs.

**Supplementary Figure 9:**
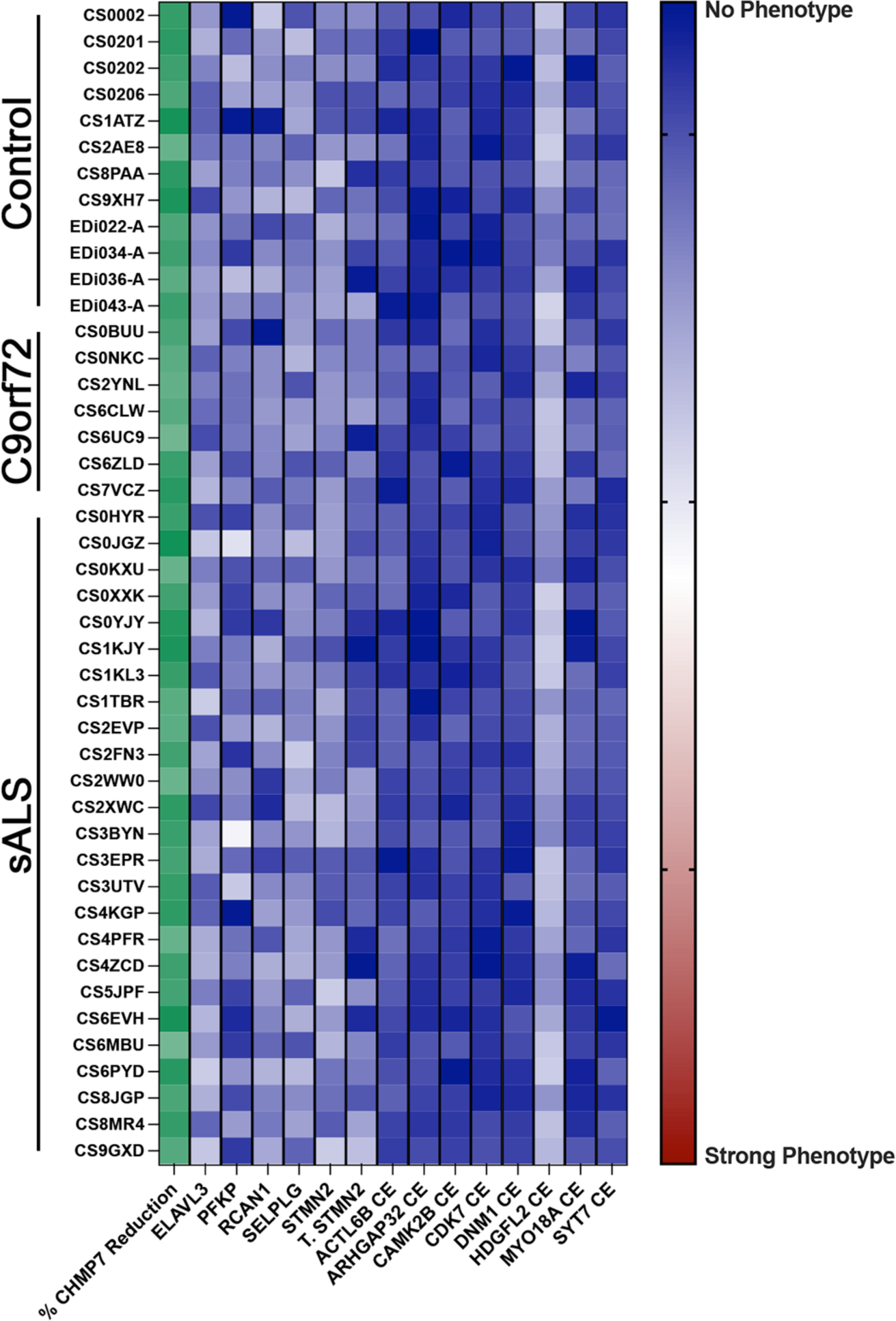
ASO mediated reduction of CHMP7 restores TDP-43 function in C9orf72 ALS/FTD and sALS patient iPSNs. Heat map depicting percentage of CHMP7 protein reduction (obtained from western blot) and TDP-43 loss of function associated alterations in target gene expression and mRNA splicing (obtained from qRT-PCR) at day 81 of differentiation following 3 weeks of treatment with 5μM CHMP7 ASO. CHMP7 ASO treatment was initiated on day 60 *following* the detection of molecular signatures of TDP-43 dysfunction. n = 12 control, 7 C9orf72, and 25 sALS iPSC lines. Boxes represent the average value across 3 independent differentiations.

**Supplementary Figure 10:**
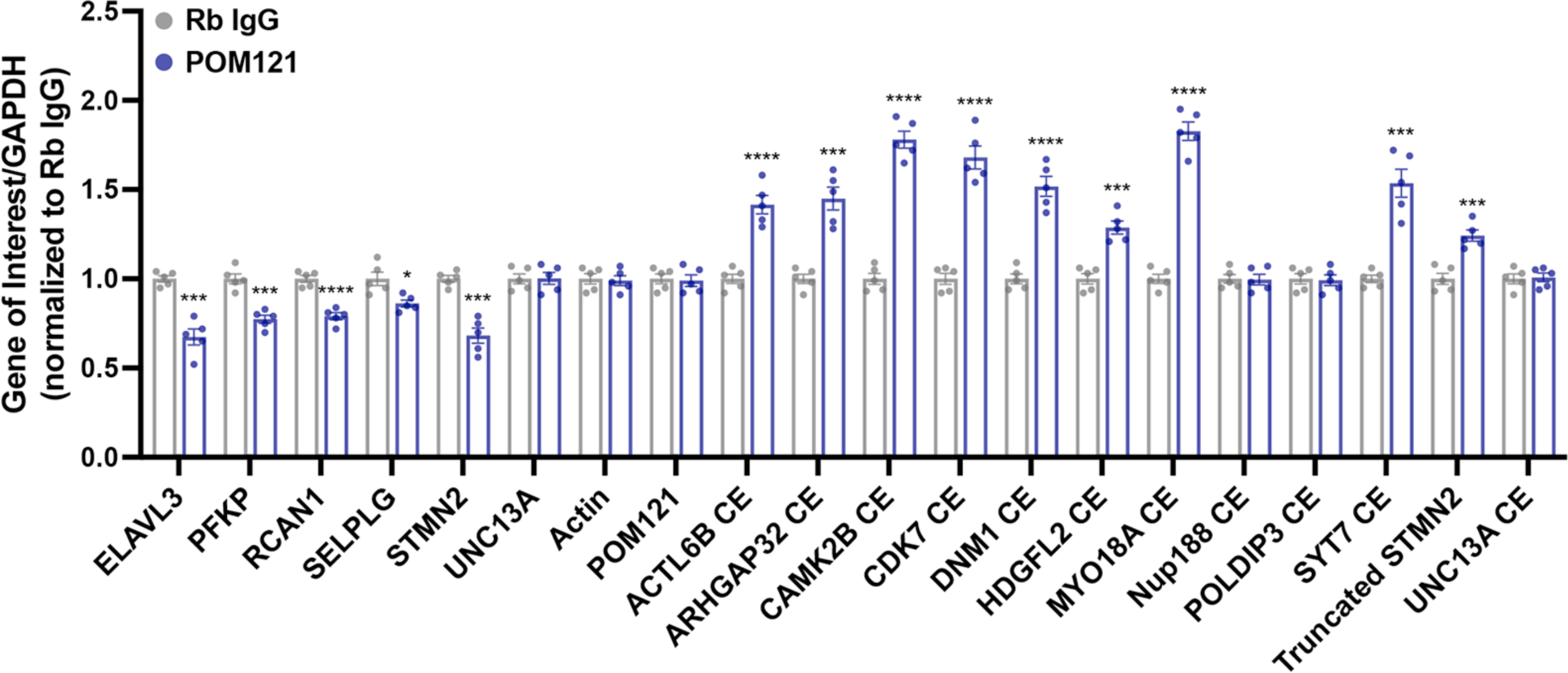
Knockdown of transmembrane Nup POM121 is sufficient to initiate TDP-43 loss of function associated changes in gene expression and mRNA splicing in wildtype iPSNs. qRT-PCR for *ELAVL3*, *PFKP*, *RCAN1*, *SELPLG, STMN2*, *UNC13A*, *ACTIN*, *POM121*, *ACTL6B* cryptic exon containing, *ARHGAP32* cryptic exon containing, *CAMK2B* cryptic exon containing, *CDK7* cryptic exon containing, *DNM1* cryptic exon containing, *HDGFL2* cryptic exon containing, *MYO18A* cryptic exon containing, *NUP188* cryptic exon containing, *POLDIP3* cryptic exon containing, *SYT7* cryptic exon containing, truncated *STMN2*, and *UNC13A* cryptic exon containing mRNA in control iPSNs 7 days following rapid degradation of endogenous POM121. n = 5 control iPSC lines. Student’s t-test was used to calculate statistical significance. *** p < 0.001, **** p < 0.0001.

**Supplementary Figure 11:**
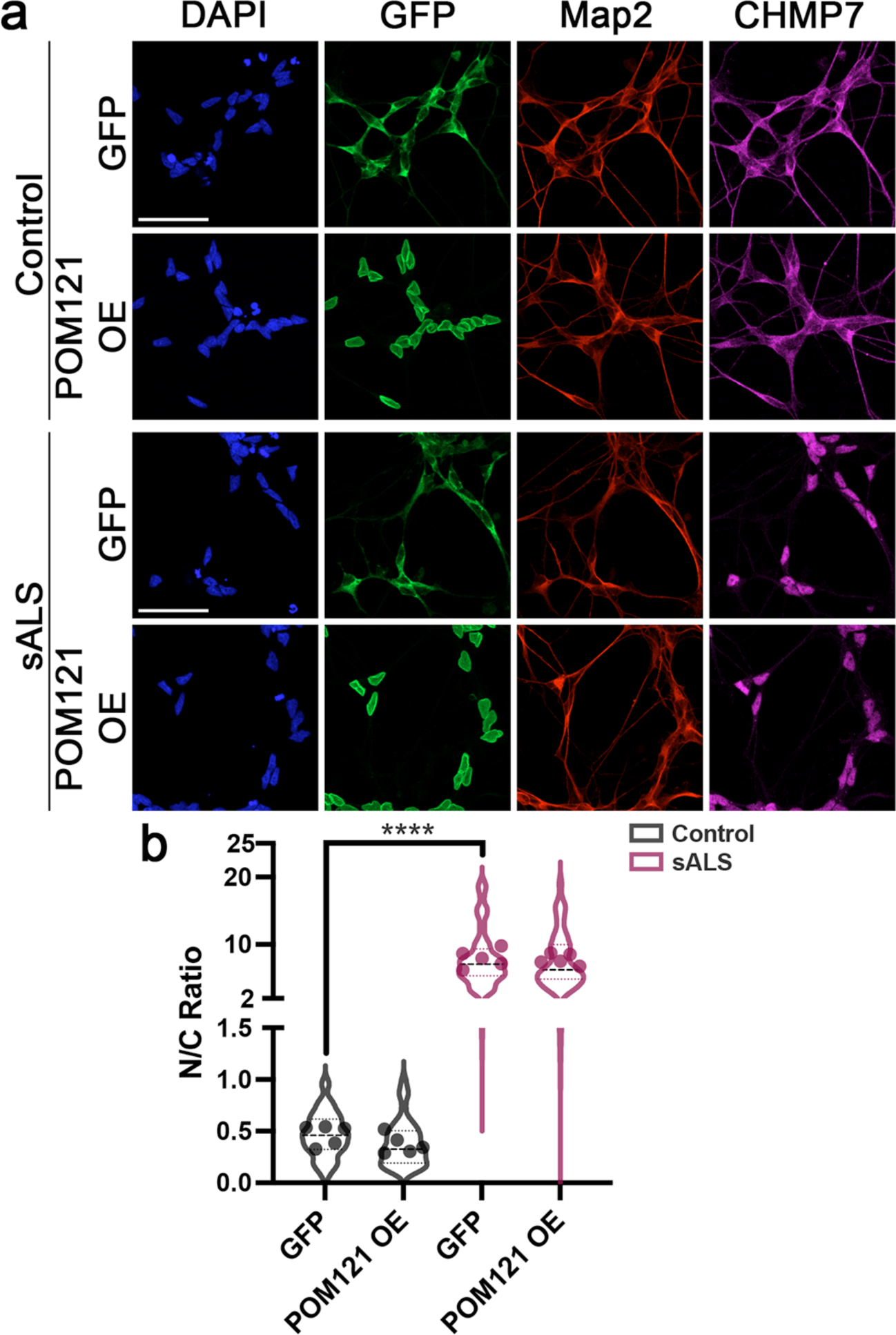
Overexpression of POM121 does not impact nuclear accumulation of CHMP7 in sALS iPSNs. **(a)** Immunostaining and confocal imaging for CHMP7 in control and sALS iPSNs following 2 weeks of GFP or POM121 overexpression. POM121 overexpression was initiated at day 46 of differentiation following the detectable initiation of TDP-43 dysfunction in sALS iPSNs. Antibody for immunostaining as indicated on top, genotype and overexpression as indicated on left. Scale bar = 50 μm. **(b)** Quantification of nuclear/cytoplasmic distribution of CHMP7. n = 5 control and 5 sALS iPSC lines, 100 Map2+ neurons per line. Two-way ANOVA with Tukey’s multiple comparison test was used to calculate statistical significance. **** p < 0.0001.

**Supplementary Figure 12:**
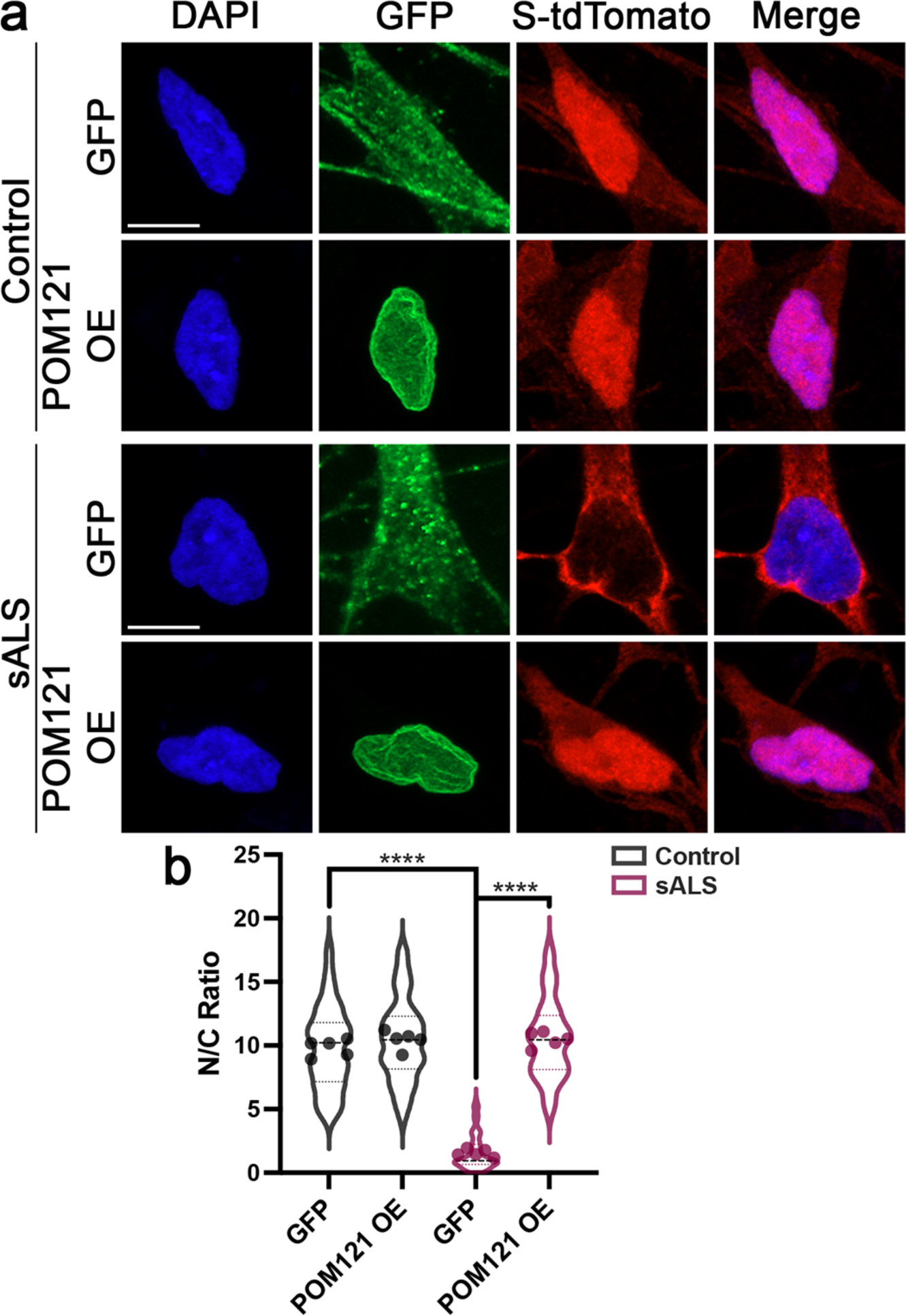
Overexpression of POM121 restores the nuclear import of the S-tdTomato NCT reporter in sALS iPSNs. **(a)** Confocal imaging for the S-tdTomato NCT reporter in control and sALS iPSNs following 2 weeks of GFP or POM121 overexpression. POM121 overexpression was initiated at day 46 of differentiation following the detectable initiation of TDP-43 dysfunction in sALS iPSNs. Stain and fluorescent protein marker as indicated on top, genotype and overexpression as indicated on left. **(b)** Quantification of nuclear/cytoplasmic distribution of the S-tdTomato NCT reporter. n = 5 control and 5 sALS iPSC lines, 100 neurons per line. Two-way ANOVA with Tukey’s multiple comparison test was used to calculate statistical significance. **** p < 0.0001.

**Supplementary Figure 13:**
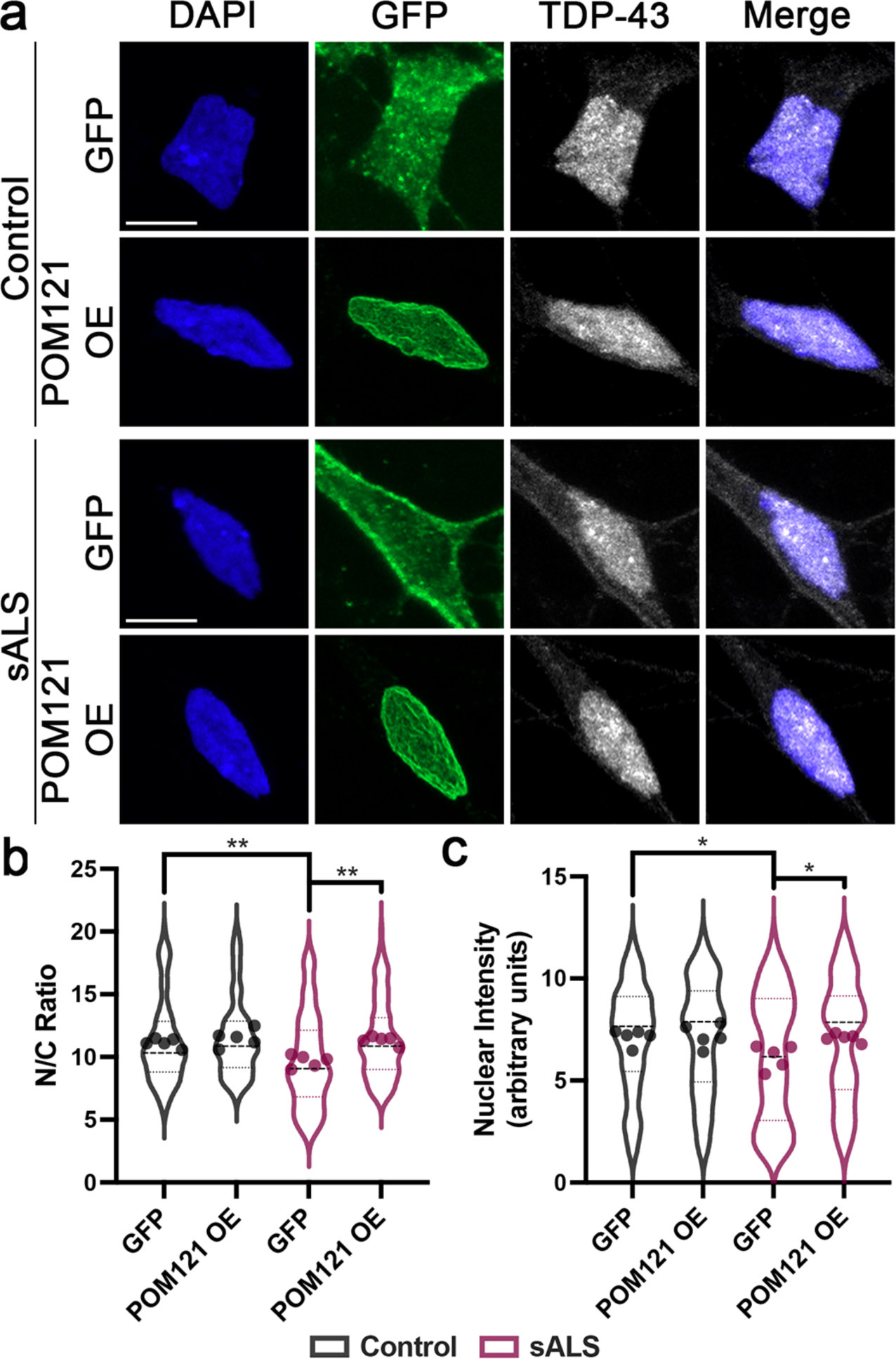
Overexpression of POM121 reverses subtle alterations in nuclear/cytoplasmic distribution of TDP-43 in sALS iPSNs. **(a)** Immunostaining and confocal imaging TDP-43 in control and sALS iPSNs following 2 weeks of GFP or POM121 overexpression. POM121 overexpression was initiated at day 46 of differentiation following the detectable initiation of TDP-43 dysfunction in sALS iPSNs. Antibody for immunostaining as indicated on top, genotype and overexpression as indicated on left. **(b)** Quantification of nuclear/cytoplasmic distribution of TDP-43. n = 5 control and 5 sALS iPSC lines, 100 Map2+ neurons per line. Two-way ANOVA with Tukey’s multiple comparison test was used to calculate statistical significance. * p < 0.05, ** p < 0.01.

